# Gamma rhythms and visual information in mouse V1 specifically modulated by somatostatin-positive neurons in reticular thalamus

**DOI:** 10.1101/2020.05.06.081877

**Authors:** Mahmood S. Hoseini, Bryan Higashikubo, Frances S. Cho, Andrew H. Chang, Alexandra Clemente-Perez, Irene Lew, Michael Stryker, Jeanne T. Paz

## Abstract

Visual perception in natural environments depends on the ability to focus on salient stimuli while ignoring distractions. This kind of selective visual attention is associated with gamma activity in the visual cortex. While the nucleus reticularis thalami (nRT) has been implicated in selective attention, its role in modulating visual perception remains unknown. Here we show that somatostatin-(SOM) but not parvalbumin-expressing (PV) neurons in the nRT preferentially project to visual thalamic nuclei. In freely behaving mice, single-unit and field recordings reveal powerful modulation of both visual information transmission and gamma activity in primary visual cortex (V1), as well as in the dorsal lateral geniculate nucleus (dLGN). These findings pinpoint the SOM neurons in nRT as powerful modulators of the visual information encoding accuracy in V1, and represent a novel circuit through which the nRT can influence representation of visual information.

## INTRODUCTION

Visual perception relies on the ability to focus on important information while ignoring distractions. Such selective attention is associated with neural oscillations in the gamma frequency band (~30-90 Hz, “gamma oscillations”) in the visual cortices in both rodents and humans (Engel et al., 2001; Taylor et al., 2005; Pavlova et al., 2006; Doesburg et al., 2008; Ray et al., 2008b; Siegel et al., 2008). Gamma oscillations, particularly in the visual cortex, are associated with a high level of cortical activity, and some have speculated that they may play a causal role in perception and the focusing of attention (Gray et al., 1992; Singer and Gray, 1995; Kreiter and Singer, 1996) or in enabling a time-division multiplexing of cortical responses to multiple simultaneous stimuli (Stryker, 1989). Electro-corticographic (ECoG) studies have shown a clear correlation between focal increases in high gamma and cortical responses to stimulation measured by other means (Ray et al., 2008a; Yazdan-Shahmorad et al., 2013). Whether gamma oscillations play a causal role in brain function or merely represent the “ringing” of an insufficiently damped response of the cortical circuit to a strong input is not clear. However, studies have shown that the effective output of primary sensory cortical areas to peripheral stimulation is much greater during a particular phase of gamma, and that increasing gamma and synchronized cortical activity by optogenetic stimulation at the appropriate phase with respect to a sensory stimulus can change the number of spikes elicited (Cardin et al., 2009; Sohal et al., 2009).

Moreover, two types of gamma oscillations have been reported in the primary visual cortex (V1), differentiated by whether they result from intra-cortical or subcortical inputs through the dorsal lateral geniculate nucleus (dLGN) (Saleem et al., 2017). The thalamus gates the flow of all visual information to the cortex. The GABAergic nucleus reticularis thalami (nRT) (Houser et al., 1980) provides a powerful source of GABAergic inhibition for thalamocortical neurons (Sherman and Koch, 1986; McCormick and Bal, 1997; Gentet and Ulrich, 2003; Lam and Sherman, 2011; Halassa and Acsády, 2016; Crabtree, 2018), allowing the nRT to control different information streams to the cortex. The nRT has been regarded as “the guardian of the gateway” to the cortex (Crick, 1984; McAlonan et al., 2008; Halassa and Acsády, 2016), and is well positioned to exert attentional selection by controlling which thalamocortical “channel” of information the cortex should “pay attention to”. Optogenetic activation of inhibitory (Gad2-positive) neurons in the dorsal portion of the nRT transiently reduces activity in the dLGN, the source of the specific sensory input to V1, particularly in its input layer (Reinhold et al., 2015). However, it is unknown whether inputs other than the strength of peripheral stimulation control the magnitude of gamma oscillations in V1.

Our previous work showed the existence of two main cell populations in the reticular thalamus: parvalbumin(PV) and somatostatin-(SOM) expressing neurons, characterized by distinct cellular and input-output circuit properties in the mouse (Clemente-Perez et al., 2017). Here, we investigated which neurons in nRT modulate activity in the thalamocortical visual system. We measured gamma power and the representation of visual information in V1 in behaving mice both during locomotion and rest, and with and without visual stimulation. We made similar measurements in dLGN. Our electrophysiological findings, in line with anatomical data, indicate that activating SOM but not PV neurons in nRT strongly reduces both visual information transmission and gamma power in V1 and dLGN.

## RESULTS

### Visual thalamocortical nuclei receive projections mainly from SOM nRT neurons

To determine whether the visual thalamocortical relay nuclei receive inputs from SOM and/or PV neurons of the nRT, we injected an AAV viral construct encoding enhanced yellow fluorescent protein (eYFP) in the nRT of SOM-Cre and PV-Cre mice. We previously validated this approach immunohistochemically and showed that SOM and PV neurons of the nRT target distinct midline and somatosensory thalamocortical relay nuclei, respectively (Clemente-Perez et al., 2017). In the nRT, SOM and PV cell bodies and their axons robustly expressed eYFP four weeks post-injection (Figure 1A). Confocal microscopy revealed dense axonal boutons from SOM nRT neurons in the dLGN, but only very sparse input from PV nRT neurons (Figure 1B-D). This was surprising given that PV neurons represent the major cellular population of nRT (Clemente-Perez et al., 2017). As previously shown, PV but not SOM neurons from the nRT projected densely to the somatosensory ventroposteromedial (VPM) thalamocortical nucleus (Figure 1B).

**Figure 1.**
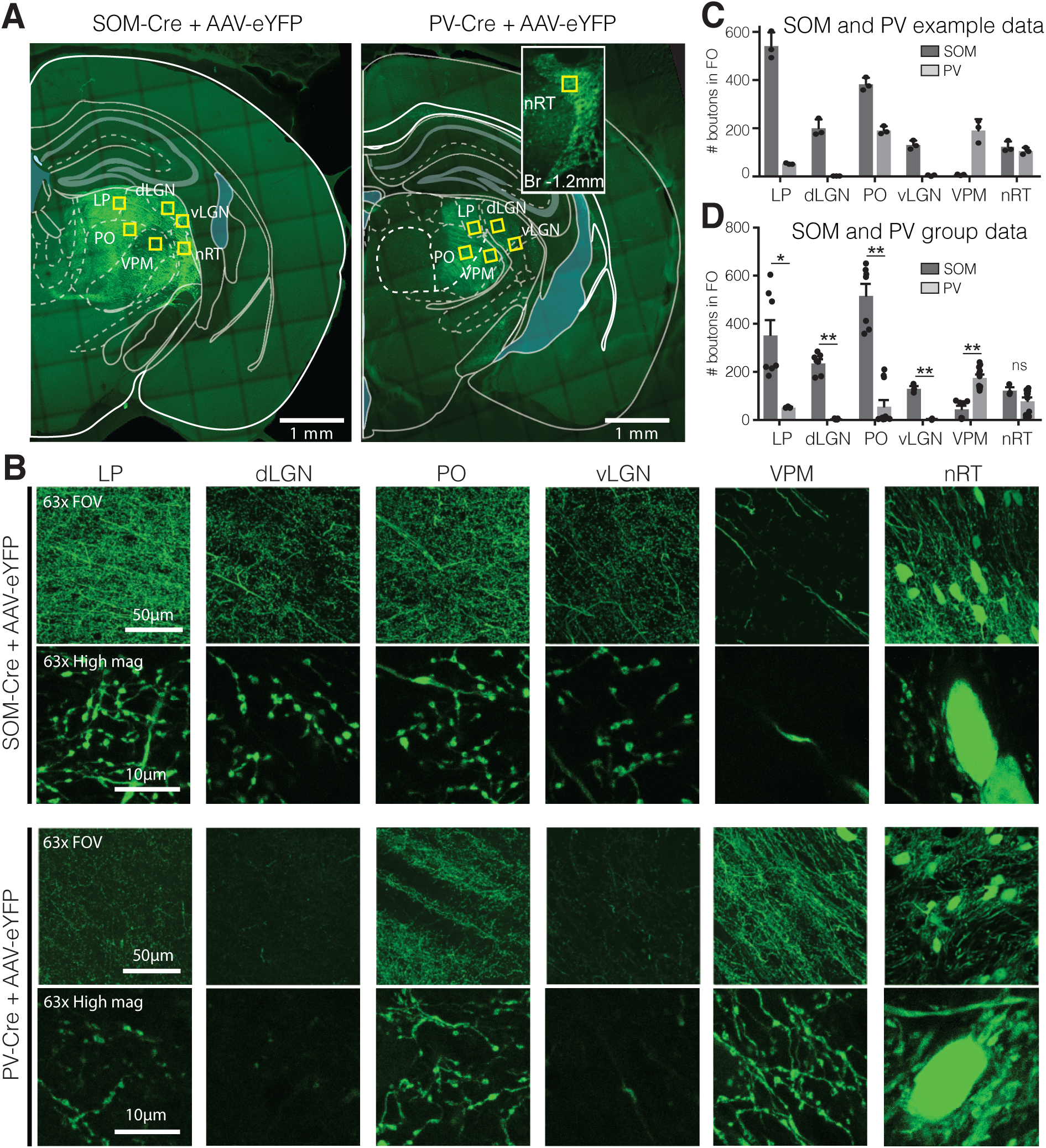
Visual relay thalamic nuclei are preferentially targeted by SOM and not PV GABAergic neurons from nRT. **A**, Representative example sections of SOM-Cre and PV-Cre mice after injection of floxed AAV in nRT, which results in eYFP expression in cell bodies and projections of SOM or PV neurons, respectively. Yellow boxes indicate locations chosen for 63x confocal imaging and putative bouton quantification. Inset: nRT injection site as seen in an adjacent section. **B**, 63x confocal images showing the entire field of view (FOV) and a zoomed cropped region (‘High mag’) to show details of axo nal boutons and nRT somata. LP: lateral posterior nucleus; dLGN: dorsal lateral geniculate nucleus; PO: posterior medial nucleus; vLGN: ventral lateral geniculate nucleus; VPM: ventroposteromedial nucleus; nRT: thalamic reticular nucleus. The expression of the viral constructs in different brain regions was confirmed using the mouse brain atlas (Paxinos and Franklin, 2001). **C**, Number of eYFP-labeled boutons present in thalamic nuclei of representative mice shown in panel A. Data taken from three consecutive sections from each mouse. **D**, Number of eYFP-labeled boutons present in thalamic nuclei of all mice imaged (n = 2 SOM-Cre, 3 PV-Cre, 3-4 sections per mouse). Differences are significant between genotypes for all regions except for nRT after correction for multiple comparisons. *p<0.05, **p<0.01.

In order to study the effect of nRT activity on the thalamocortical visual system, we transfected SOM and PV cells in the portion of the nRT that projects the visual thalamic nuclei with channelrhodopsin(ChR2) or halorhodopsin(eNpHR) containing viruses as described in (Clemente-Perez et al., 2017). This verified that the expression of opsins was located in the nRT and cell-type specific and the opsins were well ex pressed (Figure 1–figure supplement 1).

### Optogenetic activation of SOM but not PV nRT neurons reduces single-cell responses and gamma power in V1 both with and without visual stimulation

To determine whether disrupting activity of SOM nRT neurons affects visual responses in V1, we injected an AAV viral construct containing ChR2 in the nRT of SOM-Cre mice. Thereafter, extracellular recordings of single unit activity and local field potentials (LFPs) were made using a double-shank 128-channel microelectrode array placed in the V1 of mice that were free to stand or run on a polystyrene ball floating on an air stream (Figure 2A) (Du et al., 2011; Hoseini et al., 2019). Mice viewed a gray blank screen while a blue light (473 nm, ~63 mW/mm^2^) was delivered using an optical fiber implanted above nRT. Optogenetic activation of SOM cells in nRT greatly reduced the power of LFPs across almost all frequencies regardless of the locomotion state of the animal (Figure 2B). Optogenetic activation significantly reduced across-trial average power (scaled by 1/f) in all recording channels and all frequencies, especially in the gamma band (Figure 2C, Table 1). Consistent with previous findings, locomotion differently modulated power across different frequencies (Figure 2D) (Niell and Stryker, 2010). Locomotion caused a moderate power increase at low frequencies (<10 Hz), a moderate decrease in the beta band (15-30 Hz), and a dramatic power increase at higher frequencies in the gamma band (30-80 Hz) (Figure 2D and Table 1). Averaging power across all channels showed that optogenetic activation of SOM neurons in nRT nearly abolished power across the three bands in the still condition (Figure 2E, Table 1). In contrast, during locomotion optogenetic activation of SOM neurons in nRT most strongly reduced gamma power in V1 (Figure 2E, Table 1).

**Table 1.** Results of significance testing across different conditions. Power amplitudes are in units of 1,000*uV2/Hz (Figs. 2C-I, 3B-E) and firing rates are in Hz (Figs. 2K, 2L, 4F).

|  |  | <u>Still & laser off</u> | <u>Still & laser on</u> | <u>Running & laser off</u> | <u>Running & laser on</u> | <u>p-value</u> |
| --- | --- | --- | --- | --- | --- | --- |
| **Fig. 2C (n=111 channels)** | $\theta$ | 39 ± 1.8 | 27 ± 1.4 | | | 1.2e-9 |
| | $\beta$ | 64 ± 3.4 | 38 ± 2.3 | | | 4.7e-10 |
| | $\gamma$ | 131 ± 6.5 | 67 ± 4.4 | | | 2.5e-12 |
| **Fig. 2D (n=111 channels)** | $\theta$ | 39 ± 1.8 | | 45 ± 2.8 | | 0.70 |
| | $\beta$ | 64 ± 3.4 | | 49 ± 2.7 | | 4.5e-5 |
| | $\gamma$ | 131 ± 6.5 | | 201 ± 9.2 | | 5.2e-12 |
| **Fig. 2E (n=4 mice)** | $\theta$ | 29 ± 7.1 | 1.8e4 ± 5.0 | 33 ± 10.6 | 32 ± 9.5 | 0.34 (still), 0.88 (running) |
| | $\beta$ | 47 ± 10.0 | 2.7e4 ± 6.3 | 36 ± 6.0 | 32 ± 6.4 | 0.34 (still), 0.68 (running) |
| | $\gamma$ | 95 ± 18.5 | 5.0e4 ± 10.3 | 137 ± 31.04 | 83 ± 17.7 | 0.11 (still), 0.02 (running) |
| **Fig. 2H (n=94 channels )** | $\theta$ | -8.9 ± 0.5 | | 16 ± 1.1 | | 0 |
| | $\beta$ | 18 ± 1.6 | | 5.7 ± 1.1 | | 4.9e-9 |
| | $\gamma$ | 39 ± 1.4 | | -42 ± 1.6 | | 0 |
| **Fig. 2I (n=4 mice)** | $\theta$ | 6.2 ± 8.7 | 1.1e4 ± 7.7 | -0.9 ± 2.8 | 1.9e4 ± 6.4 | 0.88 (still), 0.02 (running) |
| | $\beta$ | 22 ± 11.9 | 3.3e4 ± 5.9 | 3.7 ± 2.9 | 1.5e4 ± 7.4 | 0.34 (still), 0.20 (running) |
| | $\gamma$ | 17 ± 7.6 | -2.3e4 ± 7.1 | 19 ± 8.5 | -2.8e4 ± 9.2 | 0.02 (still), 0.02 (running) |
| **Fig. 2K (n=134 BS, 51 NS)** | BS | 10.6 ± 1.61 | 6.9 ± 1.00 | 16.9 ± 2.65 | 10.5 ± .59 | 0.13 (still), 0.08 (running) |
|  | NS | 15.98 ± 2.94 | 9.2 ± 1.33 | 25.9 ± 4.34 | 15.2 ± 2.40 | 0.08 (still), 0.02 (running) |
| **Fig. 2L (n=100 BS, 32 NS cells)** | BS | -38.2 ± 0.83 | -23.6 ± 4.72 | -38.2 ± 1.01 | -34.0 ± 4.02 |  |
|  | NS | -38.5 ± 0.65 | -21.1 ± 8.10 | -38.0 ± 1.17 | -35.56 ± 5.31 |  |
| **Fig. 4B (n=121 channels)** | $\theta$ | 22 ± 1.4 | 22 ± 1.4 | | | 0.78 |
| | $\beta$ | 42 ± 5.3 | 45 ± 5.5 | | | 0.50 |
| | $\gamma$ | 77 ± 12.5 | 78 ± 12.4 | | | 0.88 |
| **Fig. 4C (n=121 channels)** | $\theta$ | 22 ± 1.4 | | 27 ± 2.0 | | 0.04 |
| | $\beta$ | 42 ± 5.3 | | 42 ± 4.5 | | .49 |
| | $\gamma$ | 77 ± 12.5 | | 99 ± 12.2 | | 0.004 |
| **Fig. 4E (59 channels)** | $\theta$ | 27 ± 0.3 | 3.3 ± 0.4 | | | 0.45 |
| | $\beta$ | 4.0 ± 0.8 | 4.7 ± 0.7 | | | 1.5e-6 |
| | $\gamma$ | 6.6 ± 1.8 | 6.5 ± 1.2 | | | 0.33 |
| **Fig. 4F (n=31 BS, 43 NS cells)** | BS | 10.8 ± 1.72 | 11.8 ± 1.81 | 15.8 ± 2.64 | 16.1 ± 2.75 | 0.66 (still), 0.96 (running) |
|  | NS | 12.0 ± 2.13 | 12.3 ± 2.15 | 18.9 ± 4.05 | 18.3 ± 3.90 | 0.85 (still), 0.92 (running) |

**Figure 2.**
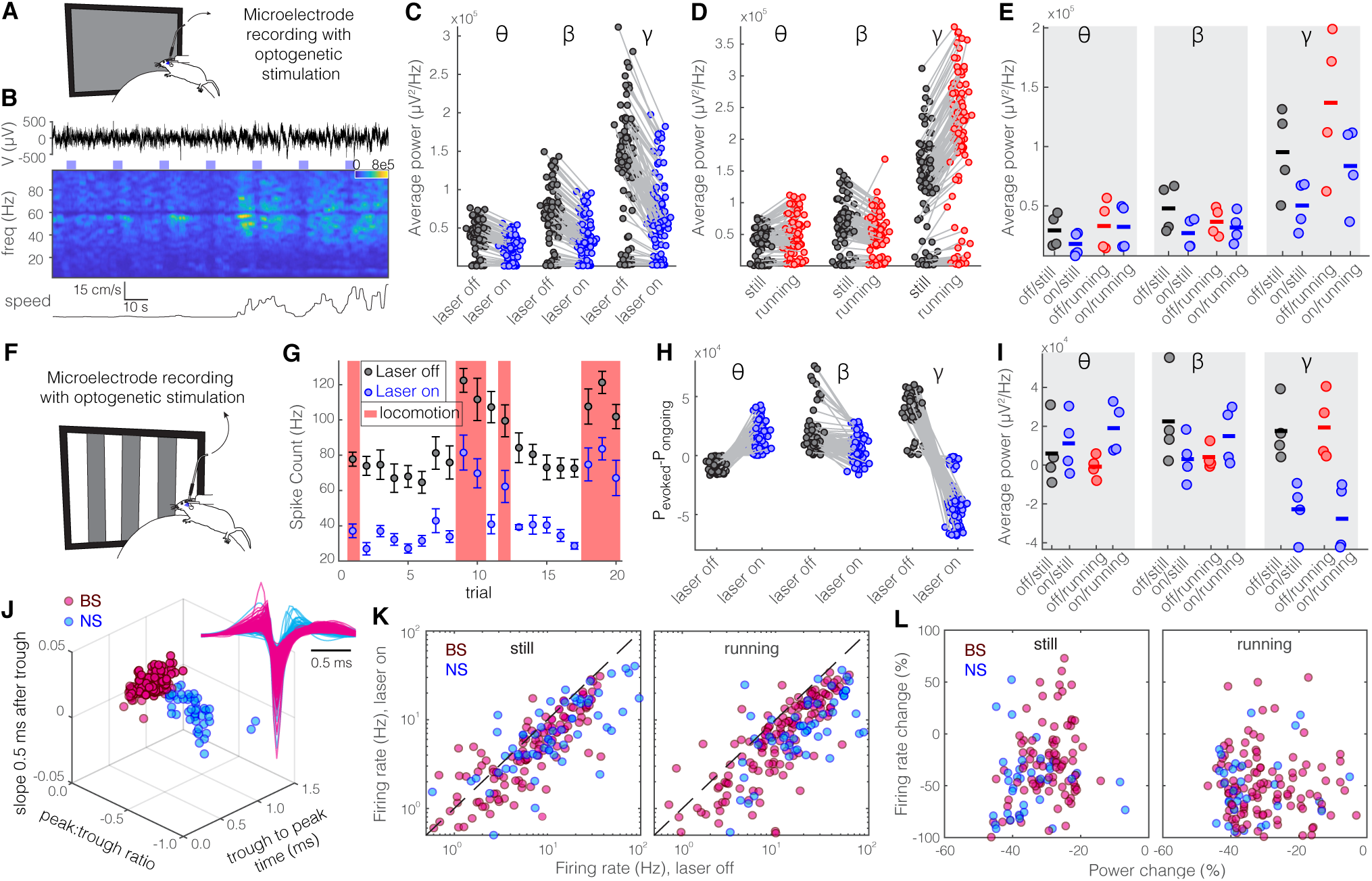
Optogenetic activation of SOM nRT neurons reduces gamma activity in the primary visual cortex (V1) both with and without visual stimulation. **A**, Neural activity was recorded from V1 in freely-moving mice. Mice were presented with a gray blank screen while a blue light (473 nm, ~63 mW/mm2) was delivered to ChR2-expressing SOM cells in nRT using an optical fiber implanted above the nRT. Mouse movement was tracked over the course of the experiment. **B**, Representative extracellular raw voltage trace is shown along with its power spectrum. Blue shading areas indicate optogenetic activation. Mouse movement speed is shown at the bottom. **C**, Across-trial average power of all channels in the absence (black) and presence (blue) of optogenetic activation in one representative mouse shows a moderate decrease across theta (4-8 Hz) and beta (15-30 Hz) bands and a strong decrease in gamma band (30-80 Hz) (See Table 1 for statistics). **D**, Across-trial average power of all channels in one representative mouse indicates that locomotion slightly modulates theta and beta powers, while causing a strong enhancement in gamma power (Table 1). **E**, Average power across all channels shows that optogenetic activation of SOM nRT cells greatly reduces power across the three bands in the still condition (black vs. blue marks). Optogenetic activation of SOM nRT cells strongly modulates gamma power when the mice are running (red vs. blue marks) (Table 1). **F**, Visual responses were recorded while mice were presented with moving gratings (8 directions, each moving in one of two possible directions; 2 s duration; randomly interleaved with optogenetic stimulation) in the visual field contralateral to the recording site. **G**, Firing rate (averaged over all drifting directions) of an example cell during the course of experiment. Black marks: visual responses when the laser is off; Blue marks: visual responses when visual stimuli and optogenetic activation of SOM nRT cells are coupled. Red shadings: locomotion state. Error bars: SEM. **H**, Stimulus evoked (average over all trials) minus ongoing power of all channels when the laser is off (black circles) versus the laser-on condition (blue circles) indicates a significant shift across all frequencies (Table 1). **I**, Same as in Fig. 2E in the presence of visual stimulus (Table 1). **J**, Using the three parameters calculated from average waveforms, cells were classified into narrow(NS, cyan) or broad(BS, magenta) spiking (height of the positive peak relative to the negative trough: −0.20 ± 0.01, −0.34 ± 0.02 (p=1.02e-9, Wilcoxon rank-sum test); the time from the negative trough to the peak: 0.73 ± 0.02, 0.32 ± 0.02 ms (p=3.9e-33), slope of the waveform 0.5 ms after the negative trough: 0.01 ± 0.00, −0.01 ± 0.00 (p=5.94e-35), BS (n=169) and NS (n=73) cells respectively). Subplot: average spike waveforms for all units, aligned to minimum, demonstrating BS (magenta) and NS (cyan) cells. **K**, Firing rate of BS (magenta) and NS (cyan) cells across different conditions (Table 1). **L**, Percentage change in firing rate of both cell types versus percentage change of power in channels that each cell is recorded from for still and running states (Table 1, Spearsman’s rho and p: 0.05, 0.58 BS & still; −0.12, 0.22 BS & running; 0.19, 0.29 NS & still; 0.05, 0.77 NS & running).

To investigate how evoked visual responses are af fected by optogenetic activation of SOM nRT cells, visual responses were recorded to drifting sinusoidal gratings presented in the visual field contralateral to the recording site, and SOM nRT neurons were activated optogenetically during randomly interleaved trials (Figure 2F). While visual stimulation selectively enhanced betaand gamma-band LFP power, optogenetic activation of SOM nRT neurons increased theta power and markedly reduced power in the other two frequency bands (Figure 2G-I–figure supplement 2, Table 1). Firing rates of isolated neurons in V1 were also modulated by optogenetic activation of SOM nRT neurons. Neurons in V1 were classified as narrowor broad-spiking using three parameters calculated from average spike waveforms (Niell and Stryker, 2008) (Figure 2J). Narrow-spiking cells (abbreviated as NS) consist of fast-spiking interneurons, whereas broad-spiking (abbreviated as BS) cells are 90% ex citatory and 10% inhibitory cells (Barthó et al., 2004; Atencio and Schreiner, 2008). Optogenetic activation of SOM nRT neurons significantly suppressed activity of both BS and NS cell types in V1 during both stationary and locomotion states (Figure 2K, Table 1). However, locomotion continued to increase the firing rates even in the presence of optogenetic activation compared with firing rates during stationary states (Figure 2K, Table 1).

To test whether different cortical layers are disproportionately modulated by the optogenetic activation of SOM nRT cells, we compared the changes in visual responses at individual recording sites. Visual stimuli evoked the strongest responses in the superficial layers (~layer 2/3) and layer (Figure 2–figure supplement 3A, B), and optogenetic activation of SOM nRT neurons reduced the activity evoked by visual stimuli in these layers (Figure 2–figure supplement 3C). Excluding the most superficial recording sites, where effects were variable, the greatest difference between evoked spike responses in the presence and absence of optogenetic activation of SOM nRT cells was between and μm from the cortical surface in every case (Figure 2–figure supplement 3D, E). Consistent with these findings, no clear relationship was observed between the changes in firing rates of individual neurons and the LFP power at their recording sites (Figure 2L).

In further experiments, we expressed ChR2 in PV nRT neurons, which constitute the major GABAergic pop ulation in this nucleus (Clemente-Perez et al., 2017). Strikingly, the effect of activating them produced completely different effects from activation of SOM nRT neurons. Activation of PV nRT cells produced consistent enhancement rather than a large reduction in spiking activity of both BS and NS cells in V1, although this effect was small (Figure 2–figure supplement 4). In addition, activating PV nRT neurons produced a small increase in gamma power during the still condition and a relatively larger increase during running, with and without visual stimulation (Figure 2–figure supplement 4).

In the absence of visual stimulation when the mouse was still, activation of SOM nRT neurons induced a change in field potential that was large in theta, and even larger in beta and gamma activity. In line with previous work (Lee et al., 2014), there was very little change in theta activity during locomotion, and only small changes in beta, but dramatic changes in gamma (Figure 3A). Activation of SOM neurons in nRT had a much more powerful effect on V1 than did activation of PV neurons. Indeed, activation of SOM nRT neurons dramatically reduced gamma whereas activation of PV nRT neurons slightly increased it whether the animal was still or running (Figure 3A). The effects of activating SOM or PV nRT cells during visual stimulation on firing rates was consistent with their effects on gamma (Figure 3B) because firing rates in BS and NS cells in V1 are known to be correlated with gamma power (for review see Wang, 2010; Buzsáki and Wang, 2012).

**Figure 3.**
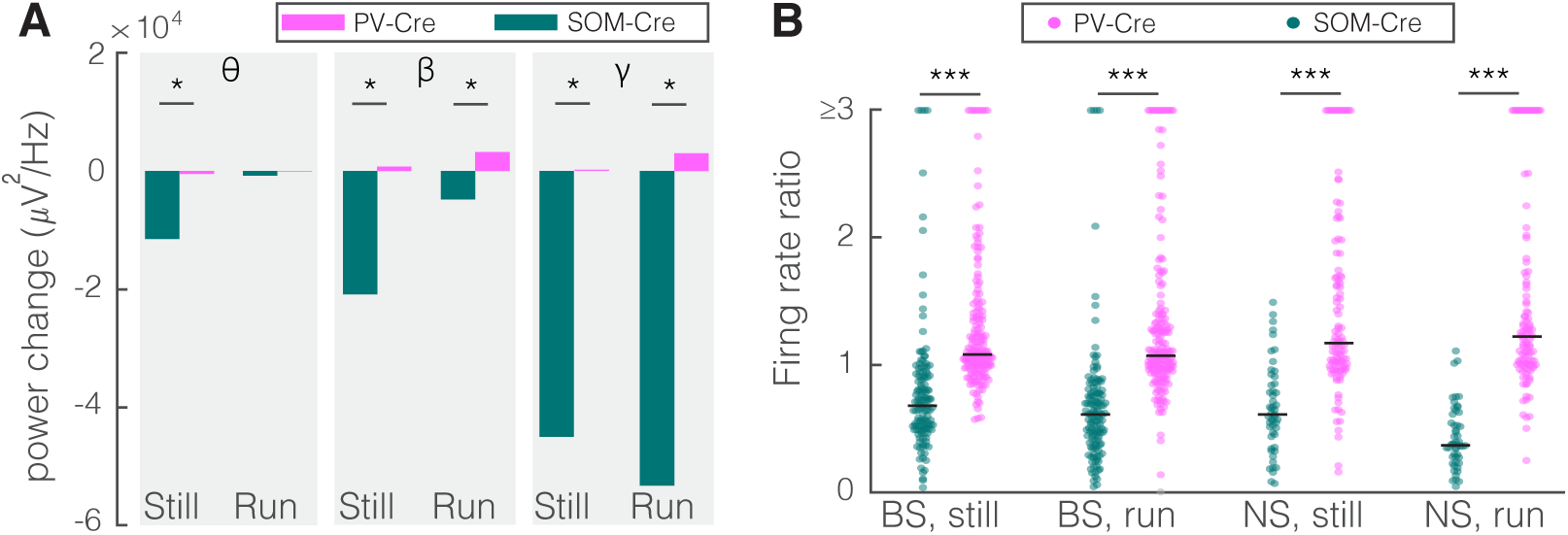
Optogenetic activation of SOM nRT neurons has much stronger effects on V1 activity than activation of PV nRT neu rons. **A**. Changes in the V1 ongoing power in response to opto genetic stimulation of SOM (teal) and PV (pink) neurons in nRT (SOM vs. PV; theta, still: −11.5e3 vs. −1.3e3, p=0.028, run: −0.7e3 vs. −12.9e3, p=0.114; beta, still: −20.8e3 vs. 4.4e3, p=0.028, run: −4.8e3 vs. 2.3e3, p=0.48; gamma, still: −45.0e3 vs. 11.9e3, p=0.028, run: −53.3e3 vs. 9.8e3, p=0.028; all powers in uV2/Hz). **B**. Effect of optogenetic activation of SOM and PV neurons in nRT on the ratio of laser on to laser off firing rates in narrow-spiking (NS) and broad_spiking (BS) cells in V1 in still and running conditions (SOM vs. PV; BS, still: 0.79 vs. 1.27, p=9e-27, run: 0.67 vs. 1.29, p=2e-31; NS, still: 0.65 vs. 1.46, p=4.7e-14, run: 1.46 vs. 1.50, p=1.6e-21). *p<0.05, **p<0.01, ***p<0.001.

As a control experiment, we tested for potential non-specific effects of the laser light used for optoge netic activation by performing recordings in the V1 cortex of SOM-Cre mice that were injected with an AAV viral construct containing eYFP in the nRT. Visual responses were recorded with and without optogenetic light during interleaved trials (Figure 4A, D). As expected, delivering the optogenetic light (blue, nm, ~63 mW/mm^2^) in nRT had no effect on the power across different frequency bands with (Figure 4E, Table 1) or without (Figure 4B, Table 1) visual stimulation; nor did it alter the firing rates of individual neurons (Figure 4F, Table 1) or the effects of locomotion on field potential powers (Figure 4C, Table 1). These results suggest that the laser light used for optogenetic experiments has no effects in the absence of opsin expression.

**Figure 4.**
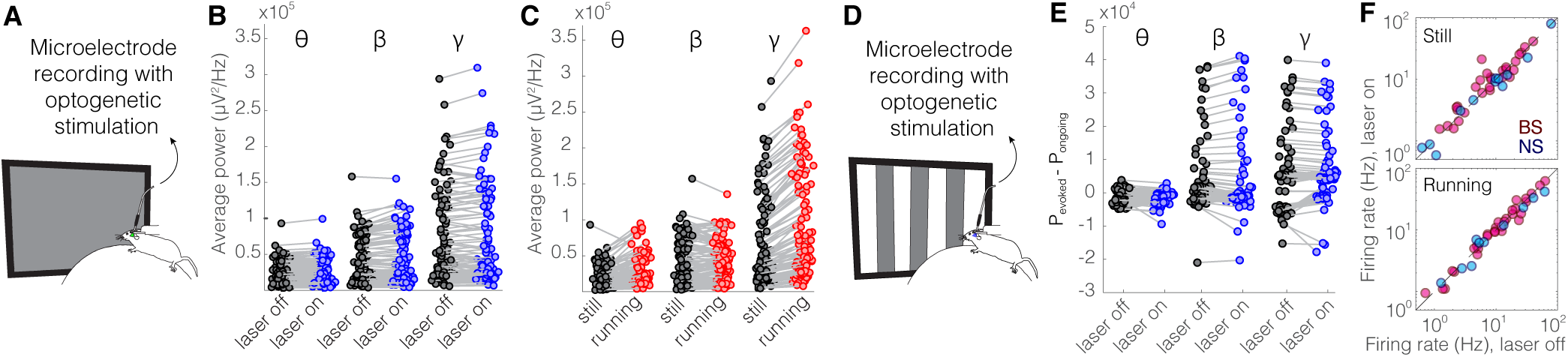
Optical stimulation of control eYFP-expressing SOM nRT neurons does not change V1 activity. **A**, Experimental setup. **B**, Across-trial average power of all channels in the absence (black) and presence (blue) of optogenetic activation in one mouse shows no change (Table 1). **C**, Across-trial average power of all channels in one mouse indicates that locomotion slightly modulates theta and beta powers, while causing a strong enhancement in gamma power (Table 1). **D**, Visual responses were recorded while mice were presented with moving gratings. **E**, Average power across all channels shows that light delivery in eYFP-expressing SOM nRT cells does not affect power in V1 (Table 1). **F**, Firing rate of BS (magenta) and NS (cyan) cells across different conditions are not affected by the light in mice in which nRT does not express the opsin (Table 1).

### Optogenetic inhibition of SOM nRT neurons enhances gamma power in V1 with and without visual stimulation

As an alternative approach to determine how nRT gates visual information, we examined the effect of inhibiting either SOM or PV nRT neurons by expressing eNpHR. The outcome of this experiment is not trivial since optogenetic activation and inactivation of neurons do not necessarily produce symmetric effects (Phillips and Hasenstaub, 2016; Moore et al., 2018). We found that inhibiting eNpHR-expressing SOM neurons produced a consistent increase in both gamma and spiking activity of cells in V1 (Figure 5–figure supplement 5). In contrast, we expressed eNpHR in PV neurons in nRT and found that inhibiting PV nRT neurons enhanced both gamma and spiking activity of cells in V1, but this effect was small (Figure 5–figure supplement 6). Notably, the effects of PV nRT inhibition on V1 were smaller than those of SOM nRT inhibition.

**Figure 5.**
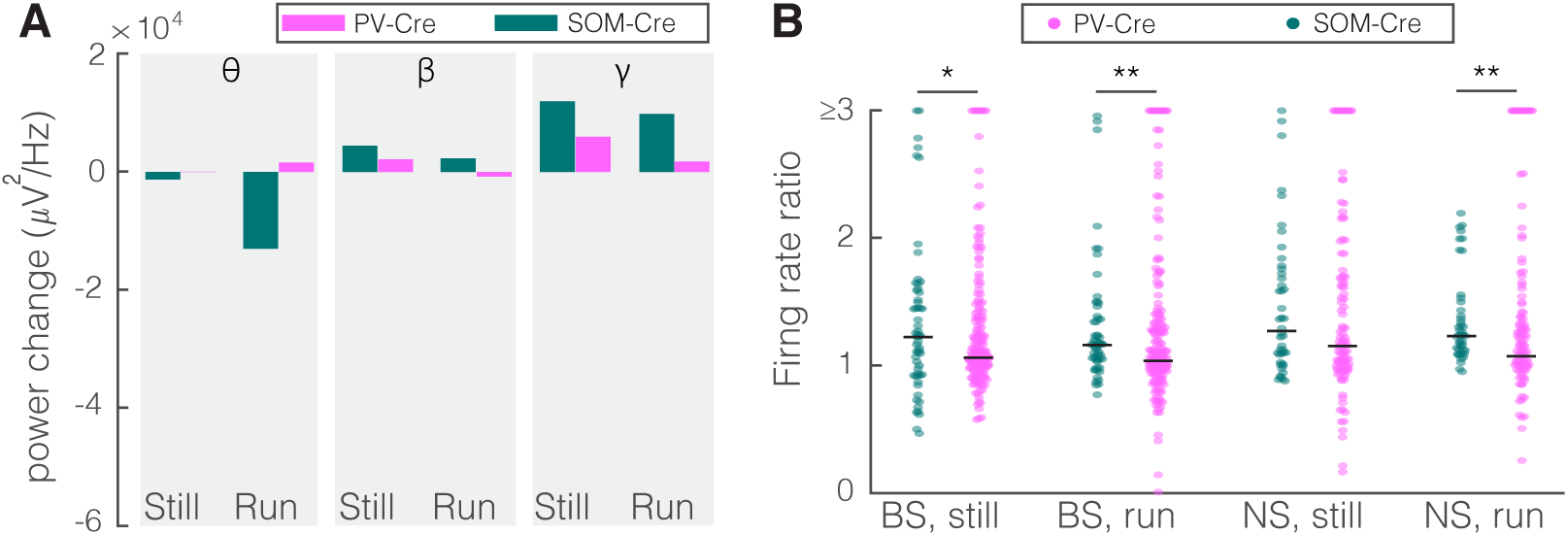
Optogenetic inhibition of SOM nRT neurons has much stronger effects on V1 activity than inhibition of PV nRT neurons on V1 activity. **A**. Changes in the V1 ongoing power due in response to optogenetic inhibition of SOM (maroon) and PV (dark yellow) neurons in nRT across the three frequency bands (SOM vs. PV; theta, still: −3.8e3 vs. 0.2e3, p=1, run: −15.9e3 vs. 3.5e3, p=0.13; beta, still: 6.4e3 vs. 4.1e3, p=0.80, run: 3.9e3 vs. −0.7e3, p=0.53; gamma, still: 11.9e3 vs. 5.4e3, p=0.4, run: 9.8e3 vs. 3.8e3, p=0.13; all powers in uV2/Hz). **B**. Effect of the optogenetic inhibition of SOM and PV neurons in nRT on the ratio of laser on to laser off firing rates in NS and BS cells in V1, in still and running conditions (SOM vs. PV; BS, still: 1.33 vs. 1.27, p=3.7e-2, run: 1.29 vs. 1.10, p=3.1e-4; NS, still: 1.47 vs. 1.26, p=5.6e-2, run: 1.35 vs. 1.17, p=5.7e-4). *p<0.05, **p<0.01, ***p<0.001.

In the absence of visual stimulation when mice were still, inhibition of SOM and PV nRT cells produced changes in field potential that were much smaller than those produced by activation of these cells and were comparable across all frequency bands (Figure 5A). During locomotion the difference between inhibiting SOM and PV nRT neurons was most striking in theta and gamma activity (Figure 5A), suggesting that the output of SOM nRT cells plays a more important role in regulating theta and gamma activity in V1 during locomotion. Consistent with the denser projection to dLGN from SOM nRT than PV nRT cells, inhibition of SOM nRT neurons during visual stimulation caused a greater increase in the firing rates of both BS and NS cells in V1 (Figure. 5B). Indeed, inhibition of PV nRT neurons caused a significant firing rate increase only during running condition and only in NS cells (Figure 5–figure supplement 6K).

### Optogenetic activation of SOM nRT but not PV nRT neurons diminishes encoding ability of BS cells in V1

The dense anatomical projection from SOM cells in nRT to visual thalamus raised the possibility that this class of cells gates the flow of visual information to V1. Since locomotion alters both strength and information content of visual responses (Niell and Stryker, 2008; Dadarlat and Stryker, 2017), we computed the mutual information conveyed by the spikes of single neurons about the visual stimuli with and without optogenetic activation of SOM nRT or PV nRT cells. Mutual information (I(R, S), see Materials and Methods) was computed separately for the running and stationary behavioral states. Optogenetic activation of SOM nRT neurons markedly reduced mutual information between the neuronal responses and our set of visual stimuli for both BS and NS cells (Figure 6A and D, Figure 6–figure supplement 7, Table 1). The reduction of mutual information in individual V1 neurons during optogenetic activation of SOM nRT cells suggests that activity of the V1 population as a whole would encode less information about visual stimuli. We estimated the representation of visual information in the response of the V1 population by training a linear decoder (LDA) to identify the visual stimulus that the animal was viewing in single stimulus trials on the basis of the spike responses in the entire population of recorded neurons. LDA incorporates the following three assumptions: that different visual stimuli evoke linearly separable responses, that evoked responses are independent across neurons, and that the responses have a Gaussian distribution. The decoder is trained on all data except a single trial which is left out for testing purposes (leave-one-out cross validation, LDA-LOOXV). This approach allows us to quantify how well orientation of the visual stimulus can be predicted for the single trials excluded from the training set.

**Figure 6.**
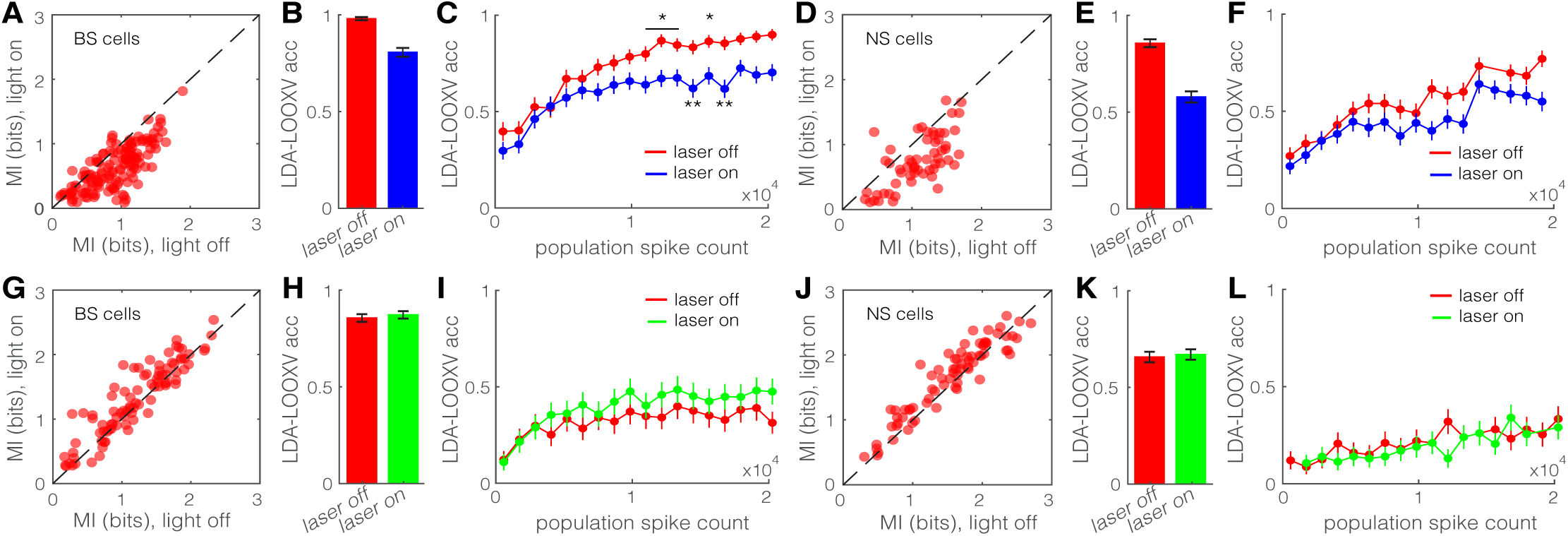
Optogenetic activation of SOM nRT neurons diminishes encoding ability in both BS and NS cells in V1. **A**, Single-cell mutual information, I(S,R), of BS neurons during locomotion for optogenetic light-off versus light-on (light-off 0.91 ± 0.03 to light-on 0.64 ± 0.03, p=2.9e-9, n=141 cells). Dashed line indicates unity. **B**, Accuracy in LDA-LOOXV classification of visual stimulus movement orientation, as a population (0.98 to 0.81, p=2e-11, Wilcoxon rank-sum test). Error bars indicate bootstrapped estimates of SE. **C**, Classification accuracy for grating movement orientation as a function of population spike count. **D**, Same as in A for NS neurons (1.09 ± 0.05 to 0.76 ± 0.05, p=2.4e-5, n=58 cells). **E**, Same as in B for NS neurons (0.86 to 0.58, p=1e-13). **F**, Same as in C for NS neurons. **G**, Same as in A during optogenetic inhibition of SOM nRT neurons (1.19 ± 0.06 to 1.31 ± 0.06, p=1.0e-4, n=90 cells). **H**, Same as in B during optogenetic inhibition of SOM nRT neurons (0.86 to 0.87, p=0.53). **I**, Same as in C during optogenetic inhibition of SOM nRT neurons. **J**, Same as in D during optogenetic inhibition of SOM nRT neurons (1.54 ± 0.07 to 1.65 ± 0.06, p=2.0e-5, n=74 cells). **K**, Same as in E during optogenetic suppression of SOM nRT neurons (0.65 to 0.67, p=0.72). **L**, Same as in F during optogenetic inhibition of SOM nRT neurons. Error bars indicate bootstrapped estimates of SE. Chance level would be at 0.16. *p<0.05, **p<0.01.

Single-trial neuronal responses during locomotion were classified less accurately during optogenetic activation of SOM nRT cells for grating orientation in both BS (98% laser-off vs. 81% laser-on, p=2e-11, Wilcoxon rank-sum test) and NS (86% vs. 58%, p=1e13) cells (Figure 6B, E). This finding indicates that the cortical representation of information about the visual world is reduced when SOM nRT cells are active, presumably because of their strong projections to the dLGN. Importantly, this finding does not depend on the behavioral state of the animal (Figure 6–Figure supplement 7C). Moreover, repeating the decoding analysis separately including only cells that are in the same range of cortical depth yielded similar accuracy (data not shown), indicating no particular laminar distinction in stimulus encoding.

Optogenetic activation of SOM nRT cells led to low er population spike counts on average, which in turn led to reduced visual information (Figure 6–Figure supplement 7A, B). The observed reduction in visu al information could therefore be due either to the re duction of neuronal firing rates or to changes in the pattern of neural responses. To distinguish between these two possibilities, we quantified decoding accuracy for trials with equal population spike counts (see Materials and Methods). We sampled (without replacement) anywhere between one and maximum number of neurons to get a very low or high number of spikes, respectively. This process was repeated 16 times and then decoding accuracy was compared for equal population spike counts during laser-off and laser-on states by including more cells in the laser-on than the laser-off classifier. LDA-LOOXV was performed separately for the data collected from each mouse and results from all four mice were pooled to generate average decoding accuracy as a function of population spike count for each cell type, behavioral state, and optogenetic state. Not surprisingly, classification accuracy increased with increasing spike count across all conditions. However, accuracy was always lower during optogenetic activation of SOM nRT cells for equal population spike counts (Figure 6C and F, Figure 6–Figure supplement 7D), and particularly so at high population spike counts. The difference was less prominent with optogenetic inhibition of SOM nRT neurons (Figure 6–Figure supplement 7E-H) and optogenetic activation or inhibition of PV nRT neurons (Figure 6–Figure supplement 8). These findings indicate that the activity of SOM nRT cells alters the pattern as well as the amount of neuronal activity in V1 that determines how accurately visual stimuli are encoded.

### Optogenetic activation of SOM nRT neurons reduces single-cell responses and gamma power in dLGN

As suggested by earlier studies (Reinhold et al., 2015), optogenetic activation of all inhibitory neurons in the nRT, regardless of the subtypes, transiently reduces activity of thalamocortical neurons in dLGN, the source of the specific sensory input to V1. But how effective is the activation of only the SOM nRT subpopulation in reducing activity in dLGN? To address this question, we expressed ChR2 in SOM nRT neurons and performed single- and multi-unit and field potential recordings from dLGN. Responses to low-frequency filtered noise stimulus were used to calculate spike-triggered averages (STAs, Figure 7A, B). STAs of less than 20% (35/193) of the recorded units display a classical center-surround structure (Figure 7B), while the rest show more complex structures (Figure 7A). Nearly all of the units were effectively inhibited during the 4-second optogenetic activation of SOM nRT neurons (**Figure 7–movie supplement 1**) and field potentials showed a dramatic reduction across all frequency bands (**Figure 7–figure supplement 9**), in agreement with a strong projection of SOM nRT neurons to dLGN (see Figure 1) (Campbell et al., 2020). While visual stimulus evoked responses in single-units (Figure 7C, top), optogenetic activation markedly reduced visual responses in single-units with a reminiscent spiking in response to the stimulus (sinusoidal drifting gratings, Hz temporal frequency, 0.04 Hz spatial frequency) (Figure 7C–E). Multi-units displayed a similar degree reduction (**Figure 7–figure supplement 9**).

**Figure 7.**
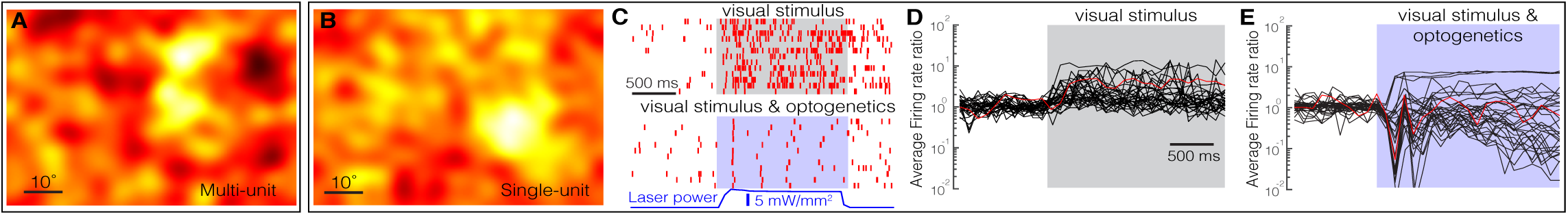
Optogenetic activation of SOM nRT neurons reduces firing in dLGN neurons. Spike-triggered averages of a multi(**A**) and a single(**B**) unit in dLGN were calculated using responses to low-frequency filtered noise stimulus. **C**, Spike raster for a representative single-unit in dLGN across several trials (rows) with visual stimulus only (top) or visual stimulus coupled with optogenetic activation of SOM nRT neurons (bottom). Grey shaded area shows the duration of the visual stimulus. Blue shading shows the duration of optogenetic activation. Firing rates of dLGN single-units in response to visual stimulation only (**D**) and in response to visual stimulation coupled with optogenetic activation of SOM nRT neurons (**E**). The firing rate was normalized to its average ongoing firing rates averaged across all stimulus conditions. Red trace in **D** and **E** is the example cell shown in **C**.

In line with our V1 recordings, optogenetic inhibition of SOM nRT neurons caused slight decrease in gamma activity and spiking in dLGN (**Figure 7–figure supplement 10**). In the absence of visual stimulation and when the animal was stationary, SOM nRT activation and inhibition showed similar changes in the amplitude of theta and gamma, but activation had a much stronger effect than inhibition on beta activity (Figure 8A). However, during locomotion, optogenetic inhibition of SOM nRT neurons had a much stronger effect than activation on theta and gamma activity, while beta activity remained more affected by activation than by inhibition (Figure 8A). During visual stimulation, which increases the firing of dLGN neurons, SOM nRT activation produced much stronger effects on dLGN firings than inhibition of SOM nRT cells (Figure 8B).

**Figure 8.**
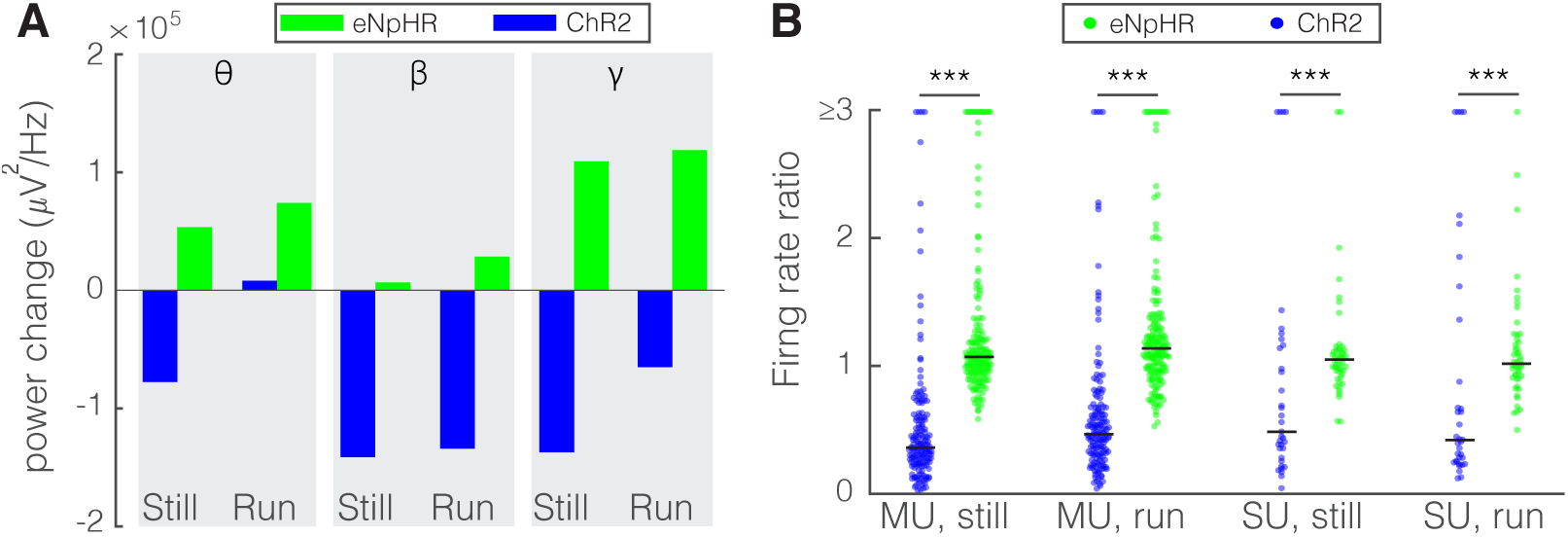
Optogenetic activation and inhibition of SOM nRT neurons bi-directionally control the activity of dLGN neurons. **A**. Changes in the dLGN ongoing power due to optogenetic activation (blue) or inhibition (green) of SOM nRT neurons across the three frequency bands (ChR2 vs. eNpHR; theta, still: −77.6e3 vs. 53.4e3, p=1, run: 8.2e3 vs. 74.1e3, p=1; beta, still: −141.3e3 vs. 6.7e3, p=0.40, run: −134.0e3 vs. 28.5e3, p=0.2; gamma, still: −137.0e3 vs. 109.3e3, p=0.4, run: −65.0e3 vs. 118.9e3, p=0.1; all powers in uV2/Hz). **B**. Effects of optogenetic activation or inhibition of SOM neurons on the ratio of laser on to laser off firing rates of dLGN cells across locomotion conditions (ChR2 vs eNpHR; MU, still: 0.65 vs. 1.28, p=5.3e-49, run: 0.66 vs. 1.65, p=7.8e-53; SU, still: 1.13 vs. 1.30, p=3.0e-12, run: 1.34 vs. 1.23, p=8.4e-12). *p<0.05, **p<0.01, ***p<0.001.

### Both activation and inhibition of SOM nRT neu rons diminish the encoding ability of single-units in dLGN

To test how nRT output alters the reliability of dLGN firing, we calculated the coefficient of variation (CV) by dividing standard deviation of firing rates in the pres ence or absence of optogenetic manipulation by their respective means. Our data reveal that optogenetic activation of SOM nRT neurons increased the CV of single-unit responses in dLGN as well as in the more numerous multi-unit responses recorded (Figure 9A). In contrast, optogenetic inhibition of SOM nRT cells slightly reduced CV of all dLGN cells (Figure 9C). These findings suggest that SOM neurons in nRT can powerfully modulate the encoding ability of neurons in dLGN.

**Figure 9.**
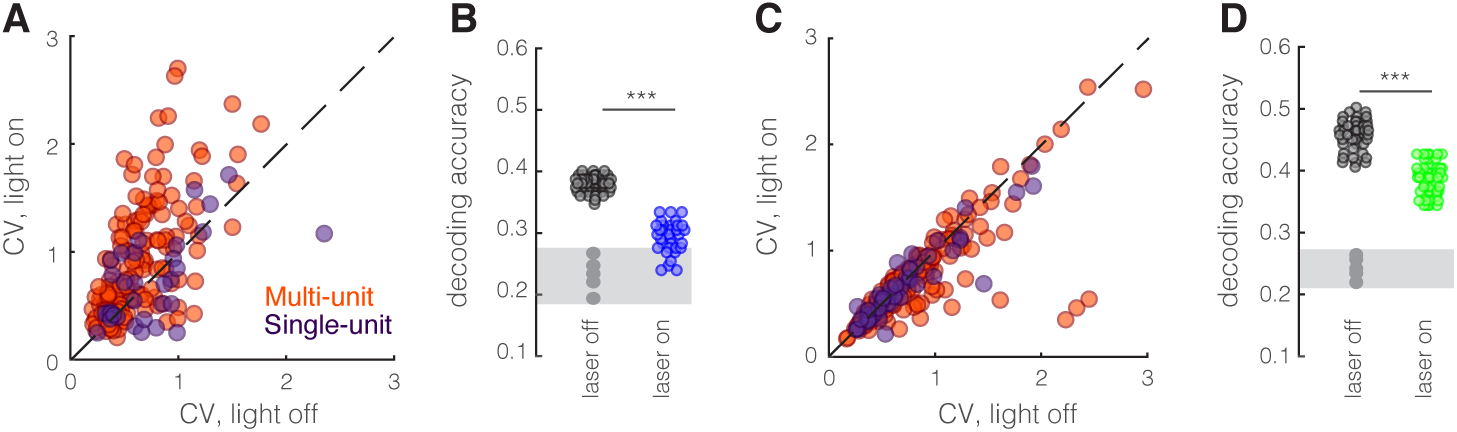
Optogenetic perturbation of SOM nRT neurons diminishes spiking reliability and stimulus location encod ing ability of dLGN cells. **A**. Coefficient of variation during optogenetic stimulation of SOM nRT neurons plotted against during visual stimulus only for multi(orange) and single(dark purple) units recorded from dLGN (multi-units: 0.65 vs. 0.96, light off vs. light on, p=1.8e-14, n=158, Wilcoxon signed-rank test; single-units: 0.75 vs. 0.87, light off vs. light on, p=0.49, n=35). **B**. Decoding of spatial location of the stimulus (see Material and Methods) is significantly less accurate in the presence of optogenetic stimulation of SOM nRT neurons (0.38 vs. 0.29, p=4.8e-15, Wilcoxon ranksum test). Grey area indicates accuracy of shuffle stimulus. **C**. Coefficient of variation during optogenetic inhibition of SOM nRT neurons plotted against during visual stimulus only for multi(orange) and single(dark purple) units recorded from dLGN (multi-units: 0.83 vs. 0.74, light off vs. light on, p=7.9e-4, n=185; single-units: 0.71 vs. 0.67, light off vs. light on, p=0.18, n=49). **D**. Decoding of spatial location of the stimulus is significantly less accurate in the presence of optogenetic inhibition of SOM nRT neurons (0.45 vs. 0.38, p=1.3e-15). Grey area indicates accuracy of shuffle stimulus.

Since dLGN neurons demonstrate relatively low orientation and direction selectivity, classification algorithms may be inaccurate. Instead, we selected ON center single-units that are not orientation or direction selective from all the recordings (n=8 units with activation, n=7 units with inhibition). Then, the responses to each stimulus was used to calculate cross-correlations between all possible pairs (28 with activation, with inhibition). Portions of the datasets were used to train deep neural networks, and the performance of the networks was evaluated on an unseen portion of data. Our results show that during optogenetic activation or inhibition of SOM neurons in the nRT, the decoding accuracy significantly diminishes (Figure 9B, D). In other words, manipulation of SOM nRT neurons blurs the representation of the visual world in the pattern of dLGN activity.

Altogether, these findings suggest that SOM nRT neurons are well positioned to control encoding ability in V1 via dLGN.

## DISCUSSION

Here we identified a new subcortical circuit that modulates both gamma power and the representation of information in the primary visual cortex. We found that the input from nRT to the dLGN in the mouse is from the class of inhibitory neurons that express soma tostatin (SOM), with negligible anatomical connections from the more numerous parvalbumin-positive (PV) neurons. Optogenetic activation of SOM nRT neurons in alert, head-fixed mice suppressed the single-unit spiking and field potential responses to visual stimulation in both dLGN and V1, and caused a second-long suppression of gamma, whether the mice were stationary or running on a polystyrene ball floating on air. Inhibition of SOM nRT cells produced mostly opposite effects. Perturbation of PV neurons in nRT had negligible effects compared to SOM nRT neurons, consistent with fewer projections to dLGN. These findings provide evidence for a specific neural circuit that regulates gamma power, and which is associated with visual attention, and the encoding of visual information in V1.

### Does nRT activity merely turn off the input to V1?

Activation of SOM nRT neurons caused a robust reduction in firing of both the thalamocortical neurons in dLGN, and the excitatory and putative fast spiking inhibitory interneurons in V1. Consistent with this effect, electron microscopy reveals geniculate terminations of SOM nRT neurons only on excitatory neurons (Campbell et al., 2020). Interestingly, activating SOM nRT neurons alters not just the amount but also the pattern of activity across the population in both dLGN and V1 leading to a blurred representation of the visual world. In contrast, despite the fact that PV cells are present in the visual portion of the nRT (see Figure 1–supple mentary figure 1), perturbation of their firing rate did not affect the visual information encoding. While optogenetic perturbation of PV nRT cells had smaller effect on V1 compared with SOM nRT cells, some of these effects were in the opposite direction. For instance, activation of PV nRT cells increased gamma activity and firing rates in V1 cells while SOM nRT neurons did the opposite. These findings are consistent with the possibility that the PV nRT neurons inhibit dLGN interneurons and/or SOM neurons, although such connections have yet to be established.

### Two pathways that modulate V1 activity

Locomotion increases both gamma power and visual responses in mouse V1, and gates a form of adult plasticity (Niell and Stryker, 2010; Kaneko and Stryker, 2014; Kaneko et al., 2017; Hoseini et al., 2019). The effects of locomotion are produced by a circuit operating through vasoactive intestinal peptide (VIP) interneurons in V1 (Fu et al., 2014, 2015). The effects of SOM nRT cells on dLGN and V1 activity were more or less additive with those of locomotion. During locomotion, when gamma power is strong in dLGN and V1, activation of SOM nRT neurons reduces it. Visual responses of both excitatory and inhibitory cortical neurons were reduced to a similar extent by activation of SOM nRT cells. These findings indicate that these two modulatory systems–locomotion via VIP cells (Niell and Stryker, 2010; Fu et al., 2014) and SOM nRT activation–contribute independently to activity in V1. In the somatosensory representation, SOM nRT cells receive inputs from mainly subcortical structures (central amygdala, anterior thalamus, external segment of globus pallidus) in contrast with PV nRT cells that mainly receive sensory cortical inputs (Clemente-Perez et al., 2017). We speculate that the SOM nRT cells are well positioned to exert a bottom-up regulation of visual attention. In contrast, the effects of locomotion on V1 activity are regulated by cortical VIP interneurons, which are ideally positioned, with dendrites in layer 1, to receive top down inputs from higher cortical areas.

### Comparison with carnivore and primate

The perigeniculate nucleus in the carnivore and primate is thought to be the portion of the nRT related to the thalamocortical visual system, although this is contested (Ahlsén et al., 1982). The carnivore perigeniculate consists of inhibitory neurons that receive excitatory input from ascending thalamocortical axons of the dLGN as well as from layer cells of V1, and they project back to inhibit the principal cells of the dLGN in a highly focused topographic fashion. This focal projection of the nRT is consistent with the Crick (1984) “Searchlight” hypothesis for nRT function. It is not known whether the nRT projection to the mouse dLGN has sufficiently precise topography to play such a focal role in directing attention to particular areas of the field. It is possible that the portion of the carnivore nRT referred to by Ahlsén et al., (1982) as “reticular neurons” may be analogous or even homologous to the nRT of the mouse. Such an arrangement could be consistent with a role for the mouse nRT in switching attention between modalities rather than among different loci in the visual field.

### Implications of our findings in disease

Sensory stimulation in the gamma range has been shown to enhance cognition in a mouse model of Alzheimer’s disease (Adaikkan et al., 2019). Given that nRT is involved in sensory processing and attention, and that its dys function has been implicated in attention disorders (Zikopoulos and Barbas, 2012; Wells et al., 2016), we propose that SOM nRT cells could be a target for modulating gamma power in V1 and visual attention.

## MATERIALS AND METHODS

### Animals

We performed all experiments in compliance with protocols approved by the Institutional Animal Care and Use Committees at the University of California, San Francisco and Gladstone Institutes. Precautions were taken to minimize stress and the number of animals used in all experiments. Adult (P60–P180) male and female mice of the following genotypes were used: SOM-Cre mice (Sst-IRES-Cre, IMSR_JAX: 018973; C57BL/6 x 129S4/SvJae); PV-Cre mice (PV-Cre, IMSR_JAX: 017320; C57BL/6 congenic); C57BL/6J mice (wild-type, IMSR_JAX: 000664).

### Viral delivery in nRT for optogenetic experiments

We performed stereotaxic injections of viruses into the nRT as described (Paz et al., 2011, 2013; Clemente-Perez et al., 2017; Ritter-Makinson et al., 2019). We targeted the nRT with the following stereotaxic coordinates: 1.3 mm posterior to bregma and −2 to −2.1 mm lateral to the midline at two different injection depths (200 nl at 2.65 and 200 nl at 3.0 mm) ventral to the cortical surface. We previously validated that this protocol results in specific expression of the viral construct in the nRT neurons and not in surrounding brain areas (Clemente-Perez et al., 2017) (see also Figure 1). To determine the effects of SOM neuron activation or inhibition on cortical rhythms and behavior, we injected AAV viruses encoding ChR2-eYFP or eNpHR3.0-eYFP in the nRT of SOM-Cre mice as previously described (Clemente-Perez et al., 2017). For control experiments, we injected AAV viruses containing eYFP in SOM-Cre mice or opsin-expressing viruses in wild-type mice. The location of viral expression was validated by histology after euthanasia in mice whose brains we were able to recover and process.

### Headplate surgery and implanting fiber optic

Three to six weeks after viral injections in the nRT, we performed a second surgery to implant a fiber optic in nRT at 1.3 mm posterior to bregma, 1.9-2.0 mm lateral to the midline and 2.3-2.5 mm ventral to the cortical surface; and a titanium headplate ‒ circular center with a mm central opening ‒ above the V1 cortex (−2.9 mm posterior to bregma, 2.5 mm lateral to the midline) or dLGN (−2.0 mm posterior to bregma, and 2.0 mm lateral to the midline). The base of the fiber optic and the entire skull, except for the region above V1 or dLGN, was covered with Metabond (Parkell Co.). One week after the recovery from this surgery, the animal was allowed to habituate to the recording setup by spending 15-30 minutes on the floating ball over the course of one to three days, during which time the animal was allowed to run freely. About two weeks following this surgery (i.e. ~4-6 weeks after viral injection in nRT), the animal’s head was fixed to a rigid crossbar above a floating ball. The polystyrene ball was constructed using two hollow 200-mm-diameter halves (Graham Sweet Studios) placed on a shallow polystyrene bowl (250 mm in diameter, 25 mm thick) with a single air inlet at the bottom. Two optical USB mice, placed mm away from the edge of the ball, were used to sense rotation of the floating ball and transmit signals to our data analysis system using custom driver software. These measurements are used to divide data into still and running trials and analyze them separately.

### Microelectrode recordings in alert mice

To control for circadian rhythms, we housed our animals using a fixed hr reversed light/dark cycle and performed recordings between roughly 11:00 AM and 6:00 PM. All the recordings were made during wakefulness in awake, head-fixed mice that were free to run on the floating ball (Figure 2A) (Hoseini et al., 2019). On the day of recording, the animal was anesthetized with isoflurane (3% induction, 1.5% maintenance) and a craniotomy of about mm in diameter was made above V1 or dLGN. After animals recovered from anesthesia for at least one hour, a 1.1-mm-long double-shank 128-channel electrode (Du et al., 2011), fabricated by the Masmanidis laboratory (University of California, Los Angeles) and assembled by the Litke laboratory (University of California, Santa Cruz), was slowly inserted through the cranial window. To record from V1, the electrode was placed at an angle of 2040° to the normal of the cortical surface and inserted to a depth of ~1000 μm. To record from dLGN, the electrode was placed at a normal angle to the cortical surface and inserted to a depth of 2.5-3.0 mm (Piscopo et al., 2013). An optical fiber (200 μm diameter) coupled to a light source (green laser for eNpHR, peak intensity ~104 mW/mm^2^ at 532 nm; blue laser for ChR2, peak intensity ~63 mW/mm^2^ at 473 nm) was connected to the implanted fiber optic in order to deliver light into nRT. Laser power (3-20mW) was measured at the end of the optical fiber before connecting to the animals. Recordings were started an hour after electrode insertion.

### Visual stimuli

Stimuli were displayed on an LCD monitor (Dell, 30×40 cm, Hz refresh rate, 32 cd/m^2^ mean luminance) placed cm from the mouse and encompassing azimuths from −10° to 70° in the contralateral visual field and elevations from −20° to +40°. In the first set of recordings, no stimulus was presented (uniform 50% gray) while nRT was exposed to the optogenetic light for s every s. For the second set of recordings, drifting sinusoidal gratings at evenly spaced directions (20 repetitions, 2 s duration, 0.04 cycles per degree, and Hz temporal frequency) were generated and presented in random sequence using the MATLAB Psychophysics Toolbox (Brainard, 1997; Kleiner et al., 2007) followed by 2-second blank period of uniform 50% gray. This stimulus set was randomly interleaved with a similar set in the presence of optogenetic light. Optogenetic stimulation was delivered for s periods beginning simultaneously with the onset of the visual stimulus, overlapping the entire stimulus period and turns off by the end of the stimulus.

### Data acquisition

Movement signals from the optical mice were acquired in an event-driven mode at up to Hz, and inte grated at 100-ms-long intervals and then converted to the net physical displacement of the top surface of the ball. A threshold was calculated individually for each experiment (1-3 cm/s), depending on the noise levels of the mouse tracker, and if the average speed of each trial fell above the threshold, the mouse was said to be running in that trial. Running speed of the animal was used to divide trials into running and still states that were analyzed separately. Data acquisition was performed using an Intan Technologies RHD2000-Series Amplifier Evaluation System, sampled at kHz; recording was triggered by a TTL pulse at the moment visual stimulation began. Spike responses during a 1000 ms period beginning 500 ms after stimulus onset were used for analysis.

### Single-neuron analysis

The data acquired using 128-site microelectrodes were sorted using MountainSort (Chung et al., 2017), which allows for fully automated spike sorting and runs at 2x real time. Manual curation after a run on minutes of data takes an additional minutes, typically yielding (range –130) isolated single units. Using average waveforms of isolated single units recorded from V1, three parameters were defined in order to classify single units into narrow- or broad-spiking (Niell and Stryker, 2008). The parameters were as follows: the height of the positive peak relative to the negative trough, the slope of the waveform 0.5 ms after the negative trough, and the time from the negative trough to the peak (see Figure 2J). For dLGN recordings, spike triggered averages (STAs) were used to classify units into singleand multi-units (Figure 7A, B).

### Mutual information

Neuronal responses are considered informative if they are unexpected. For example, in the context of visually evoked neural activity, if a neuron responds strongly to only a very specific stimulus, e.g. photographs of Jennifer Aniston (Quiroga et al., 2005), the response is informative. In contrast, if a neuron consistently produces a similar number of spikes per second to all presented stimuli, this response provides little information. This notion can be formalized by a measure of information called the Shannon entropy,

where H(X) is in units of bits. The neuron that responds to Jennifer Aniston’s face has high entropy and is therefore said to be informative. The concept is further extended to mutual information,, which quantifies how much information the neuronal response (R) carries about the visual stimulus (S) by computing the average reduction in uncertainty (entropy) about the visual stimulus produced by observing neuronal responses. Intuitively, observing responses from the aforementioned Jennifer Aniston neuron leaves little uncertainty as to which face was presented. Mutual information between S and R is calculated as follows:

where r and s are particular instances from the set of neural responses (measured as spike counts) and stimuli (grating movement directions), respectively. We used Information Theory Toolbox in MATLAB to compute mutual information (https://www.mathworks.com/matlabcentral/fileexchange/35625-information-theory-toolbox)

### Population-based analysis: decoding the visual stimulus from population responses

Data trials were separated into equal numbers of laser-off and -on trials. We then randomly subsampled from each 50 times to get a distribution of decoding errors based on the data included. We trained a linear discriminant analysis (LDA) classifier to classify single-trial neural responses, assuming independence between neurons (a diagonal covariance matrix). We used a leave-one-out approach to train and test classification separately for each condition (LDA-LOOXV). The classifier was trained and tested using MATLAB’s fitcdiscr and predict functions. To decode only grating orientation and not movement direction, we grouped stimuli moving 180° apart into the same class.

### Population-based analysis: decoding with equal population spike counts

To determine whether optogenetic manipulation of firing rates are the sole determinants of changes in information content of neuronal responses, we compared decoding accuracy from trials in laser-off and laser-on conditions with equal population spike counts, the sum of spikes from all neurons. This was accomplished by selecting subsets of neurons from the population (1-70 neurons were randomly sub-sampled with replacement). The constructed datasets retain higher-order structure between neural activity with the population but have many samples of laser-off and laser-on trials with the same population spike counts. We used an LDA-LOOXV to train and test classification separately for each subset. For each number of neurons, we subsampled with replacement 100 times from the population, yielding 100 combinations of neurons. Classifiers were trained separately on each subsample and for each condition.

### Decoding stimulus orientation in dLGN recordings

Since most dLGN units are not orientation or direction selective, predicting stimulus orientation using the LDA-LOOXV approach is not appropriate. Instead, we decided to use patterns of correlations in single-unit responses to decode stimulus identity. First, we selected all the on-center single-units that were not orientation selective from all the recordings in each condition. Using these non-simultaneously recorded neurons, we computed correlations among all possible pairs (28 pairs with optogenetic inhibition and with optogenetic activation of SOM nRT neurons) across roughly randomly selected trials for each stimulus orientation. Then the data was divided into sets of training (80%), cross-validation (10%) and testing (10%). A deep neural network was trained on the training set and its hyperparameters were tuned using its performance on the cross-validation dataset. Finally, the network predictions were tested using the test set. This process was repeated for each condition. A complete description of the model and codes are available at: https://github.com/Mahmood-Hoseini/Decoding-stimulus-orientation-in-dLGN-recordings

### Immunostaining, Microscopy, and Image Analysis

Immunohistochemistry on mouse brain sections and image analysis were performed as previously described (Clemente-Perez et al., 2017; Ritter-Makinson et al., 2019). Briefly, serial coronal sections (50 μm thick) were cut on a Leica SM2000R Sliding Microtome and immunostained with antibodies against parvalbumin (host: rabbit; concentration: 1:500; Swant catalog # PV27) and donkey anti-rabbit Alexa Fluor (concentration: 1:500; Abcam catalog # ab150076). Free-floating sections were washed in PBS, permeabilized with 0.5% Triton-X100 in PBS (PBST-0.5%), washed with 0.05% Triton-X100 in PBS (PBST-0.05%), and blocked in 10% normal donkey serum (Jackson Immunoresearch #017-000-121) in PBST-0.05% for 1 hour at room temperature. Primary incubation was performed in 3% normal donkey serum overnight at 4°C, and followed by 3×10 min washes in PBST-0.05%. Secondary incubation was performed in 3% normal donkey serum overnight for 1-2 hours at room temperature, followed by washes in PBST-0.05% and PBS.

Sections were mounted in an antifade medium (Vectashield; from Vector Laboratories, H-1000) and imaged using either a Biorevo BZ-9000 Keyence microscope or a confocal microscope. The expression of the viral constructs in different brain regions was confirmed with reference to two standard mouse brain atlases (Paxinos and Franklin, 2001) and the Allen Brain Atlas (Lein et al., 2007).

### Experimental design and statistical analysis

The experiments reported here were designed to determine (i) whether specific cell types in nRT project to visual thalamus and (ii) whether optogenetic activation or inhibition of either SOM or PV cells in nRT alters visual responses in V1 and dLGN. For the quantification of putative synaptic inputs from SOM and PV nRT neurons to various thalamocortical nuclei (see Figure 1), we performed immunohistochemistry of brain sections from n=2 SOM-Cre and n=3 PV-Cre mice (male, age range 2 to 4 months) in which we had injected an AAV construct that resulted in eYFP expression in SOM or PV neurons, respectively (for specific details, see Figure 1). For the physiology experiments, we recorded V1 responses of n=4 (2 males and 2 females) SOM-Cre mice with optogenetic activation, n=4 (2 males and 2 females) SOM-Cre mice with optogenetic inhibition, n=4 (3 males and 1 female) PV-Cre mice with optogenetic activation, and n=2 (males) PV-Cre mice with optogenetic inhibition (age range 3 to 5 months). Furthermore, we recorded dLGN responses in n=3 (males) SOM-Cre mice with optogenetic activation and n=3 (males) SOM-Cre mice with optogenetic inhibition. For the control experiment, we recorded n=2 mice (males, age 3 months) which were injected with a viral construct expressing eYFP in nRT. The expression and specific location of the opsins was verified in all the recorded mice listed above (see representative examples in Figure 1–Supplemental Figure 1). All data are illustrated in Figures 1–9 and Supplementary Figures 1–10. The main sources of variability for all of optogenetic experiments are the individual neurons and recording sites.

All numerical values are given as means ± SEM and error bars are SEM unless stated otherwise in the figure legends. Parametric and non-parametric tests were chosen as appropriate and were reported in Table 1 and figure legends. Data analysis was done in MATLAB (SCR_001622), Origin 9.0 (Microcal Software, SCR_002815), GraphPad Prism 6 (SCR_002798), R-project (SCR_001905), and SigmaPlot (SCR_003210) using Wilcoxon rank-sum, Wilcoxon signed-rank test, Spearman rank-order correlation with the Bonferroni correction for multiple comparisons (*p<0.05, **p<0.01, ***p<0.001).

## AUTHOR CONTRIBUTIONS

Conceptualization, M.S.H., M.P.S., J.T.P.; Opsin injections in nRT, J.T.P., F.S.C., A.C.P., B.H.; Surgical implantation of optical fibers and headplate implants for unit V1 recordings: M.S.H., J.T.P., B.H.; Optogenetic stimulations and recordings *in vivo*: M.S.H.; Histology and imaging: A.H.C., I.L., F.S.C.; Data Analysis, M.S.H.; Writing - Original Draft, M.S.H., M.P.S. and J.T.P.; Writing - Review and Editing, M.S.H., M.P.S., and J.T.P.; Funding Acquisition, B.H., M.P.S. and J.T.P.

## ACKNOWLEDGEMENTS

J.T.P. is supported by NIH/NINDS grantR01NS096369, Gladstone Institutes, the Kavli Institute for Fundamental Neuroscience, Michael Prize, DoD (EP150038), and NSF award #1608236. M.S.H. is supported by NSF award 1822598 and by NIH grant R01EY025174. B.H. is supported by the American Epilepsy Society Postdoctoral Research Fellowship. M.P.S. is supported by NIH grants R01EY002874 and R01EY025174. M.P.S. is a recipient of the Disney Award for Amblyopia Research from Research to Prevent Blindness. A.H.C. is supported by the Berkelhammer award. F.S.C is supported by the National Science Foundation Graduate Research Fellowship and the National Research Service Award Fellowship (NRSA, NINDS). A.C.P. is supported by National Science Foundation Graduate Research Fellowship awards #1650113. This work was also supported by an NIH/NCRR grant (C06 RR018928) to Gladstone Institutes. We thank Marie Burkart and Stephanie Holden for assistance with immunohistology, and Meredith Calvert of Gladstone Histology & Light Microscopy Core for help with confocal microscopy. We also thank Kathryn Claiborn for critical feedback on our manuscript.

---

**Figure 1–figure supplement S1.**
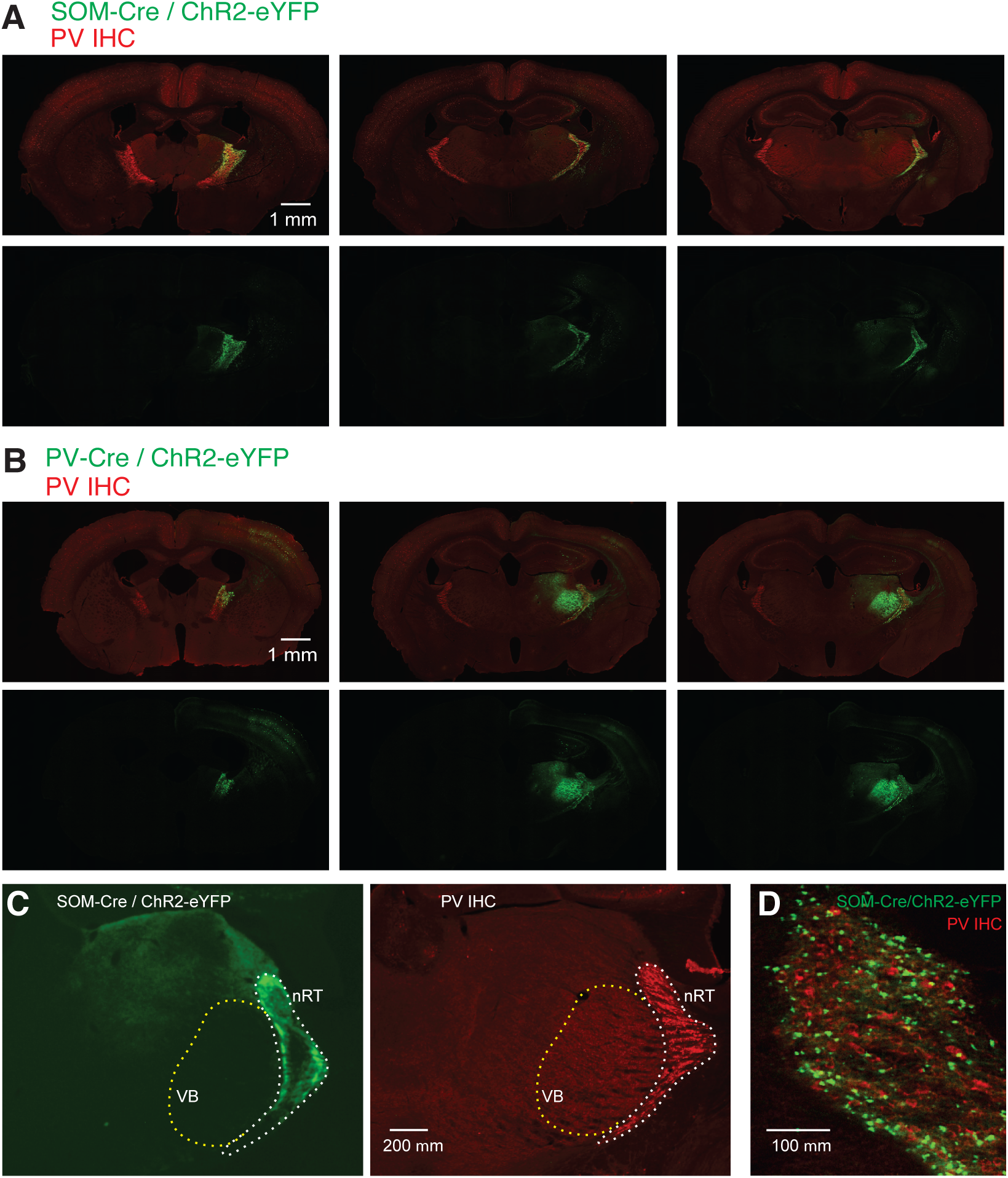
Opsin expression in SOM-Cre and PV-Cre mice. **A**, Representative composite images (individual images, 10×) of coronal brain sections from SOM-Cre mice labeled with PV-antibody (red). The sections span anterior, middle, and posterior portions of the nRT and were obtained 4-6 weeks after an intra-nRT injection of an AAV virus containing ChR2-eYFP targeting SOM-Cre cells in nRT. **B**, same as A but for PV-Cre mice. **C**, Representative composite images (individual images, 10×) of coronal brain sections from SOM-Cre mice labeled with PV-antibody (red). The sections were obtained 4-6 weeks after an intra-nRT injection of an AAV virus containing ChR2-eYFP targeting SOM-Cre cells in nRT. Note dense axonal projections from PV nRT cells to the somatosensory ventrobasal thalamus (VB), but not to dLGN, and the opposite for SOM nRT cells. **D**, A high magnification image showing that both PV (red) and SOM (green) neurons are present in the visual sector of the nRT. The results are representative of at least six mice from each group.

**Figure 2–figure supplement 2.**
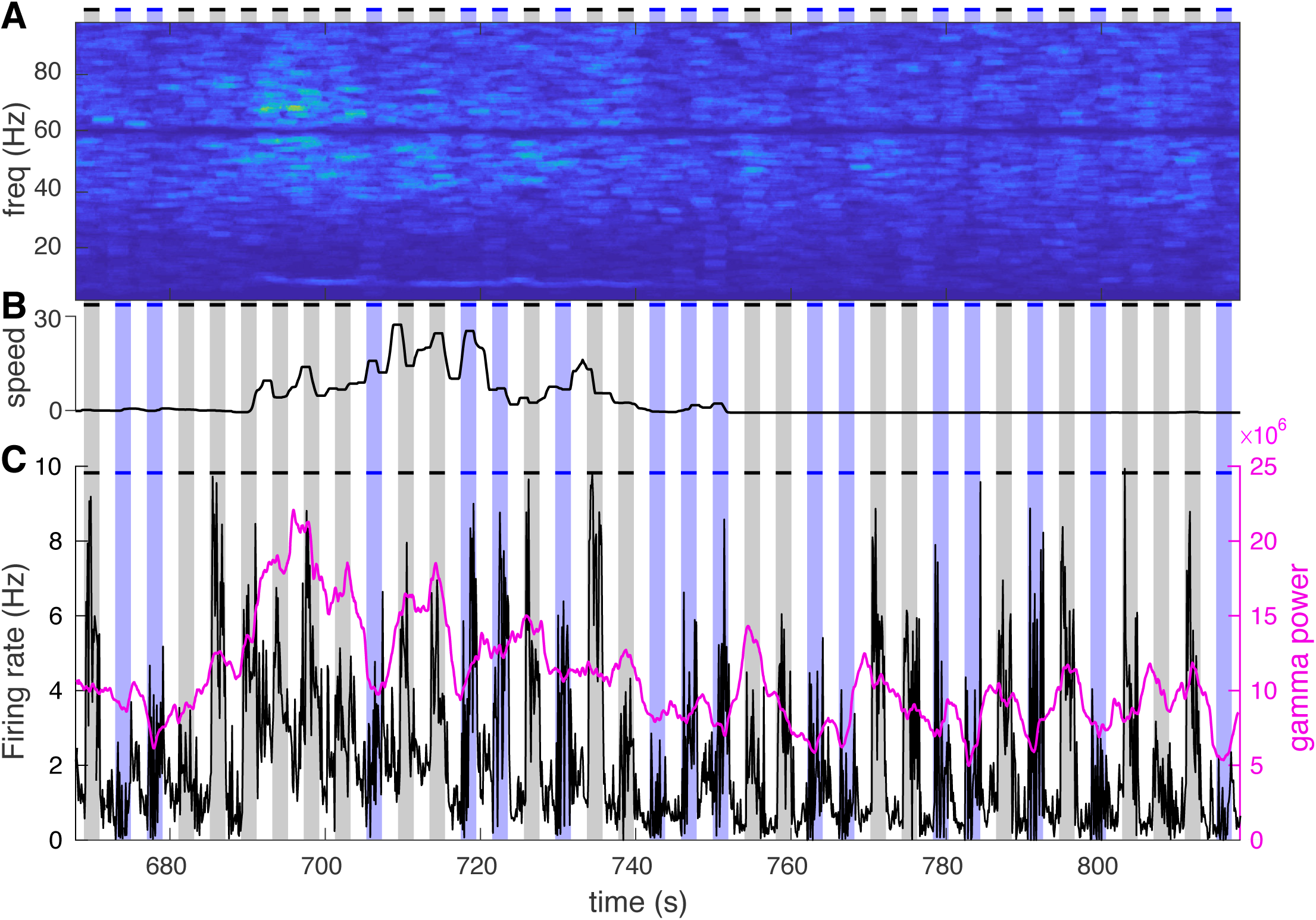
Optogenetic activation of SOM nRT neurons reduces gamma activity in the V1 cortex during visual tasks. **A**, Representative power spectrum during 150 s of visual exposure only (gray shadings) and visual exposure coupled with optogenetic activation of nRT neurons (blue shadings). **B**, Mouse movement speed. **C**, Average firing rate of all cells (black curve) is shown along with the gamma power (magenta curve). Visual stimuli consistently increase both power and average firing rate while visual exposure coupled with optogenetic activation causes a marked reduction in both measures.

**Figure 2–figure supplement 3.**
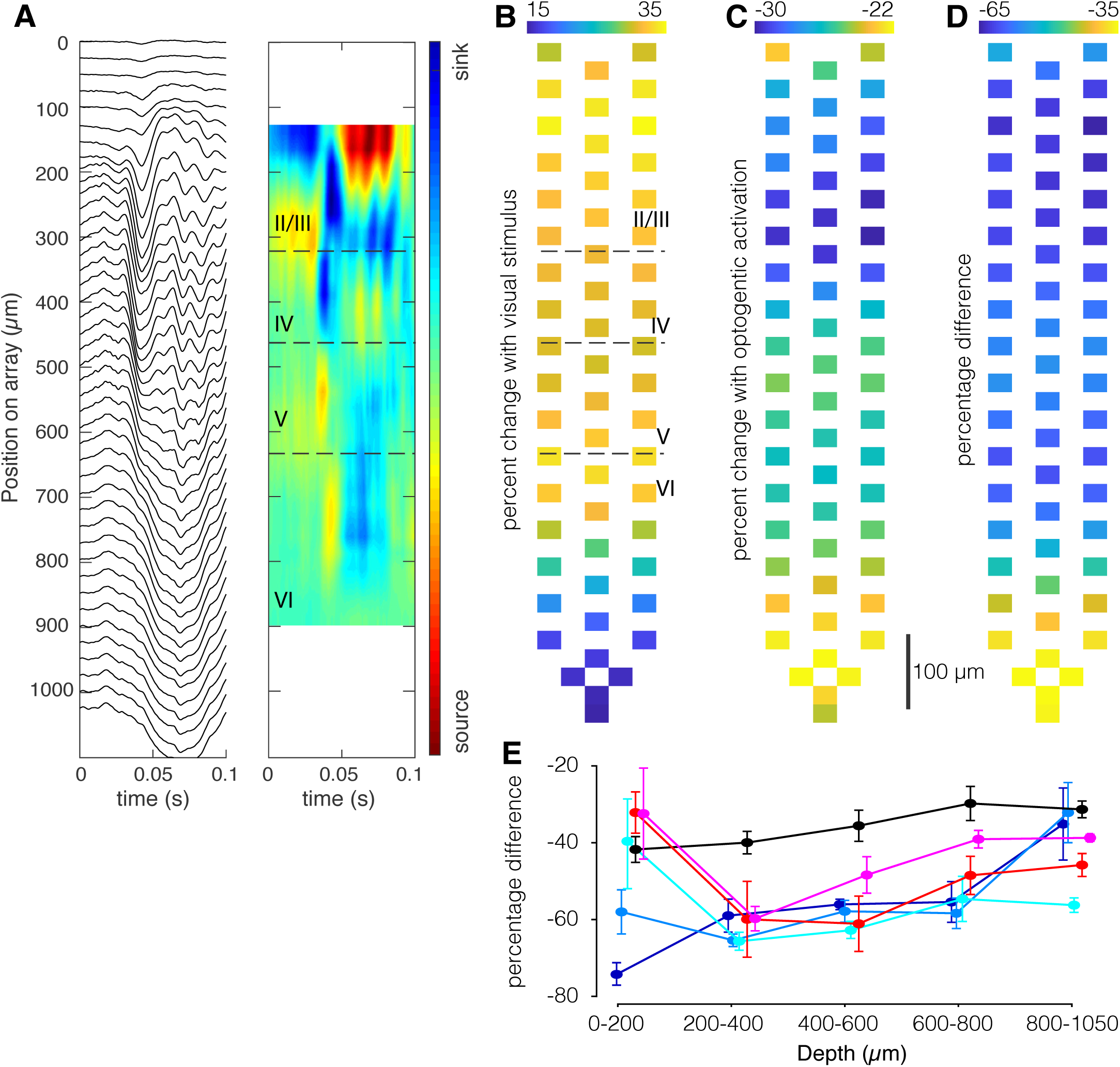
Optogenetic activation of SOM neurons in nRT has a stronger effect on neuronal responses in superficial layers in V1. **A**, Current source density identifies laminar structure in V1. Local field potential traces averaged over 150 repetition of a contrast-reversing square checkerboard stimulus (left) and current-source density plot for one mouse show laminar structure (right). Distances at left refer to electrode location relative to the top-most recording site of the array. **B**, Percentage increase of average evoked visual responses relative to the average preceding ongoing activity overlaid over microelectrode map shows an increase of about 15 to 35%. **C**, Percentage change of average evoked visual activity in the presence of optogenetic light relative to the average preceding activity overlaid over microelectrode map. **D**, Percentage difference between evoked responses in the presence of laser light (panel C) and evoked responses in the absence of laser light (panel B). **E**, Percentage difference (panel D) quantified in bins of microns across shanks in recordings from 4 mice.

**Figure 2–figure supplement 4.**
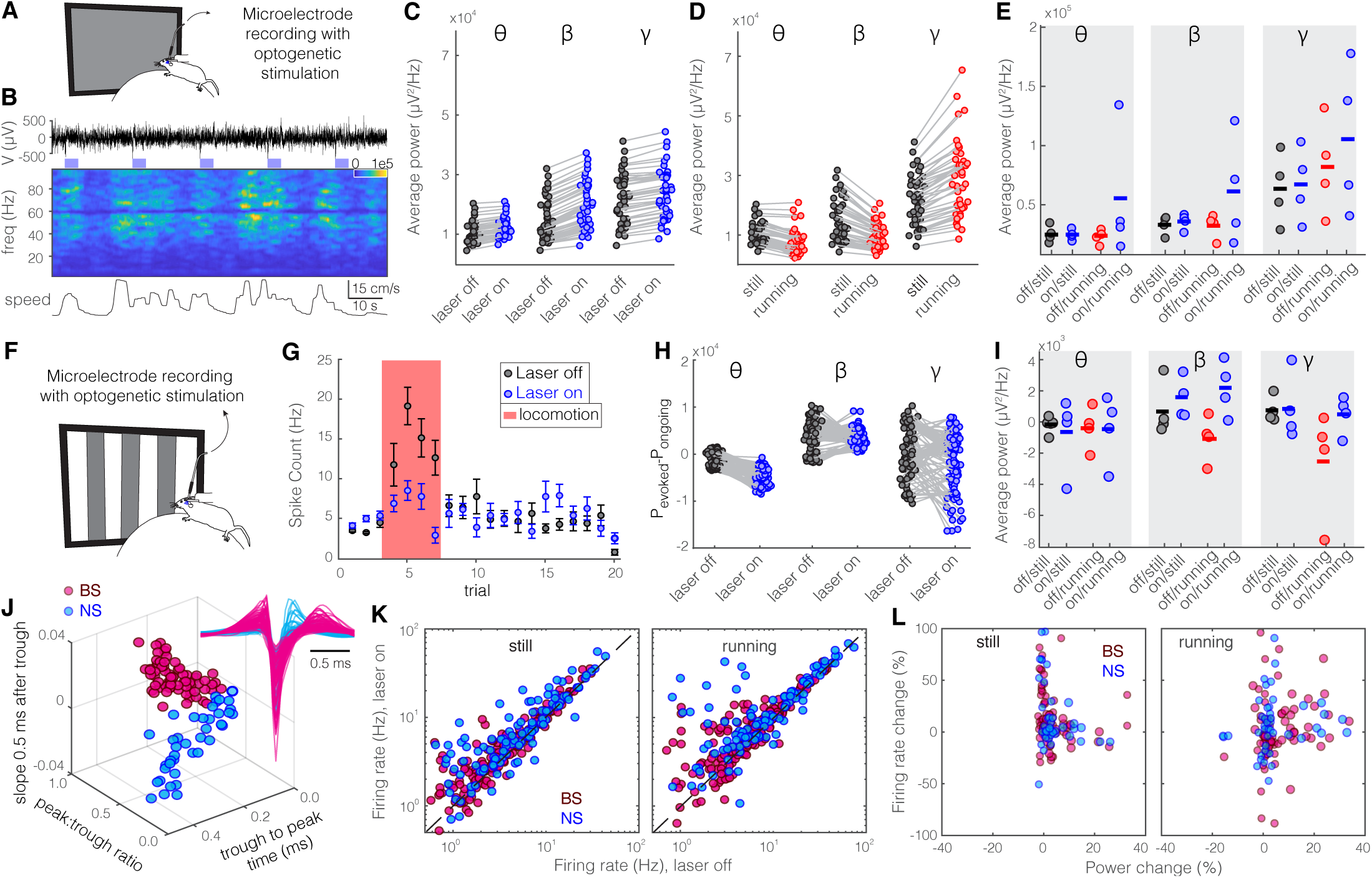
Optogenetic activation of PV nRT neurons increases gamma activity in V1 both with and without visual stimulation. Same layout as Figure 2. Here are the statistics: **C**, theta: 10.4e3 vs. 11.9e3, p=1.1e-7 Wilcoxon-signed rank test; beta: 14.6e3 vs. 19.4e3, p=1.1e-7; gamma: 21.8e3 vs. 24.1e3, p=1.1e-7. **D**, theta: 10.4e3 vs. 7.4e3, p=1.4e-8; beta: 14.6e3 vs. 9.7e3, p=1.1e-8; gamma: 21.8e3 vs. 29.2e3, p=1.1e-7. **E**, theta: −0.2e3 still & off vs. −0.7e3 still & on, p=0.68, −0.4e3 running & off vs. −0.4e3 running & on, p=0.88; beta: 0.7e3 still & off vs. 1.5e3 still & on, p=0.34, −1.0e3 running & off vs. 2.2e3 running & on, p=0.06; gamma: 0.8e3 still & off vs. 0.9e3 still & on, p=0.88, −2.5e3 running & off vs. 0.5e3 running & on, p=0.11. **I**, theta: 18.1e3 still & off vs. 17.2e3 still & on, p=1, 15.7e3 running & off vs. 47.8e3 running & on, p=0.34; beta: 26.1e3 still & off vs. 28.9e3 still & on, p=0.68, 25.2e3 running & off vs. 55.1e3 running & on, p=0.48; gamma: 57.5e3 still & off vs. 61.4e3 still & on, p=0.68, 76.6e3 running & off vs. 101.2e3 running & on, p=0.68.

**Figure 5–figure supplement 5.**
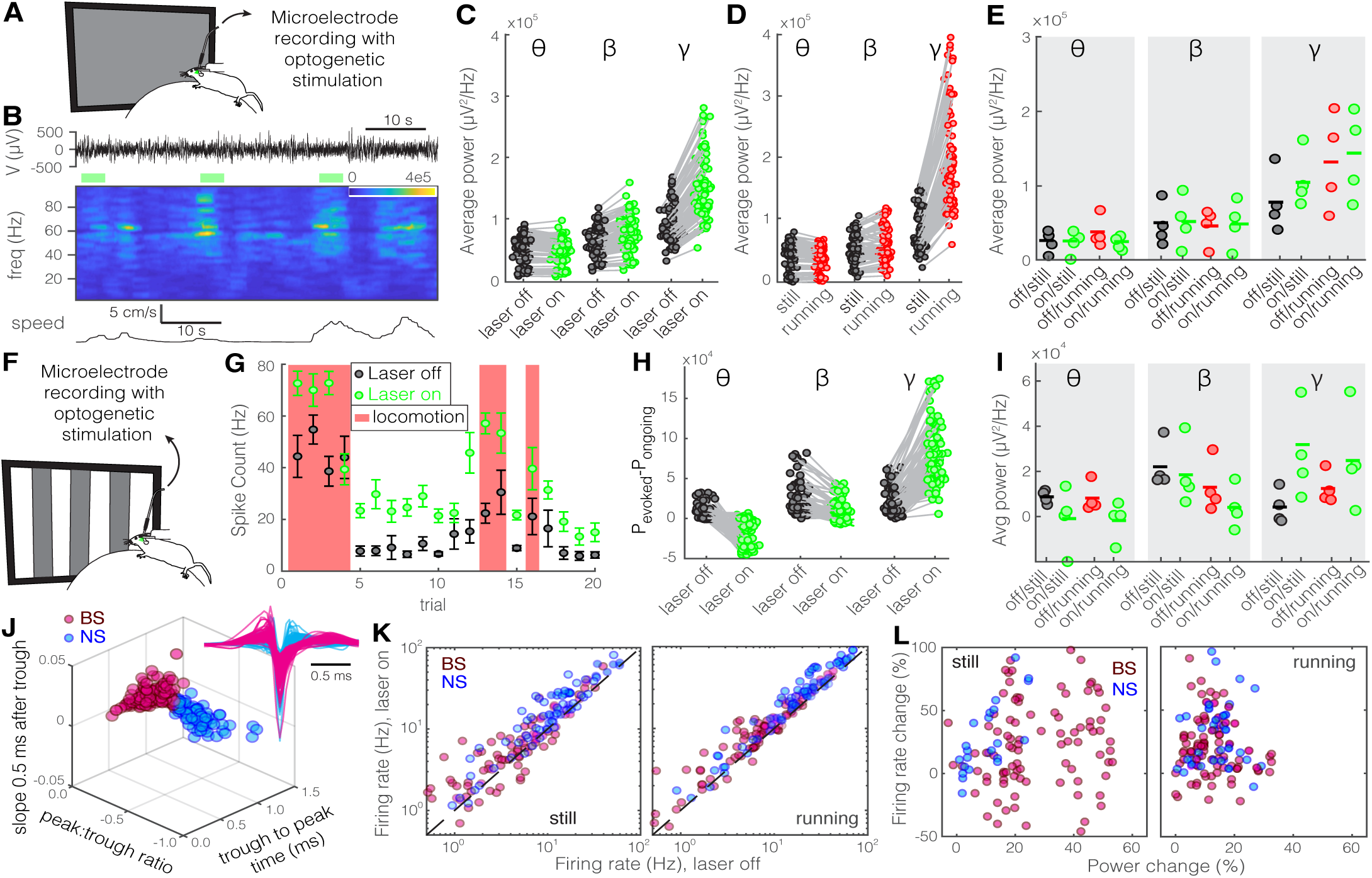
Optogenetic inhibition of SOM nRT neurons increases gamma activity in V1 both with and without visual stimulation. Same format as Figure 2. Here are the statistics: **C**, theta: 54e3 vs. 49e3, p=9.2e-3, Wilcoxon-signed rank test; beta: 63e3 vs. 77e3, p=5.2e-5; gamma: 90e3 vs. 146e3, p=0. **D**, theta: 54e3 vs. 49e3, p=0.01; beta: 63e3 vs. 68e3, p=0.48; gamma: 90e3 vs. 197e3, p=0. **E**, theta: 45e3 still & off vs. 44e3 still & on, p=0.143, 56e3 running & off vs. 43e3 running & on, p=0.34; beta: 66e3 still & off vs. 70e3 still & on, p=0.88, 65e3 running & off vs. 67e3 running & on, p=0.88; gamma: 94e3 still & off vs. 119e3 still & on, p=0.48, 130e3 running & off vs. 169e3 running & on, p=0.48. **H**, theta: 6e3 vs. −21e3, p=0; beta: 20e3 vs. 7e3, p=2.4e-10; gamma: 16e3 vs. 74e3, p=0. **I**, theta: 8.7e3 still & off vs. −0.9e3 still & on, p=0.34, 8.2e3 running & off vs. −2.0e3 running & on, p=0.11; beta: 22e3 still & off vs. 18e3 still & on, p=0.34, 13e3 running & off vs. 4e3 running & on, p=0.34; gamma: 4.0e3 still & off vs. 32e3 still & on, p=0.05, 12e3 running & off vs. 25e3 running & on, p=0.48. **J**, Height of the positive peak relative to the negative trough: BS −0.19 ± 0.01 vs. NS −0.28 ± 0.01 (p=3.2e-8, Wilcoxon rank-sum test); the time from the negative trough to the peak: 0.74 ± 0.01 vs. 0.34 ± 0.01 ms (p=5.9e-30); slope of the waveform 0.5 ms after the negative trough: 0.01 ± 0.00 vs. −0.01 ± 0.00 (p=1.2e-27); (n=97 BS and n=79 NS cells). **K**, BS & still: 8.0 (Hz) laser off vs. 9.8 laser on, p=0.002; BS & running: 11.3 laser off vs. 13.4 laser on, p=3.0e-7; NS & still: 20.4 laser off vs. 29.5 laser on, p=1.1e-6; NS & running: 31.6 laser off vs. 41.9 laser on, p=2.1e-8.

**Figure 5–figure supplement 6.**
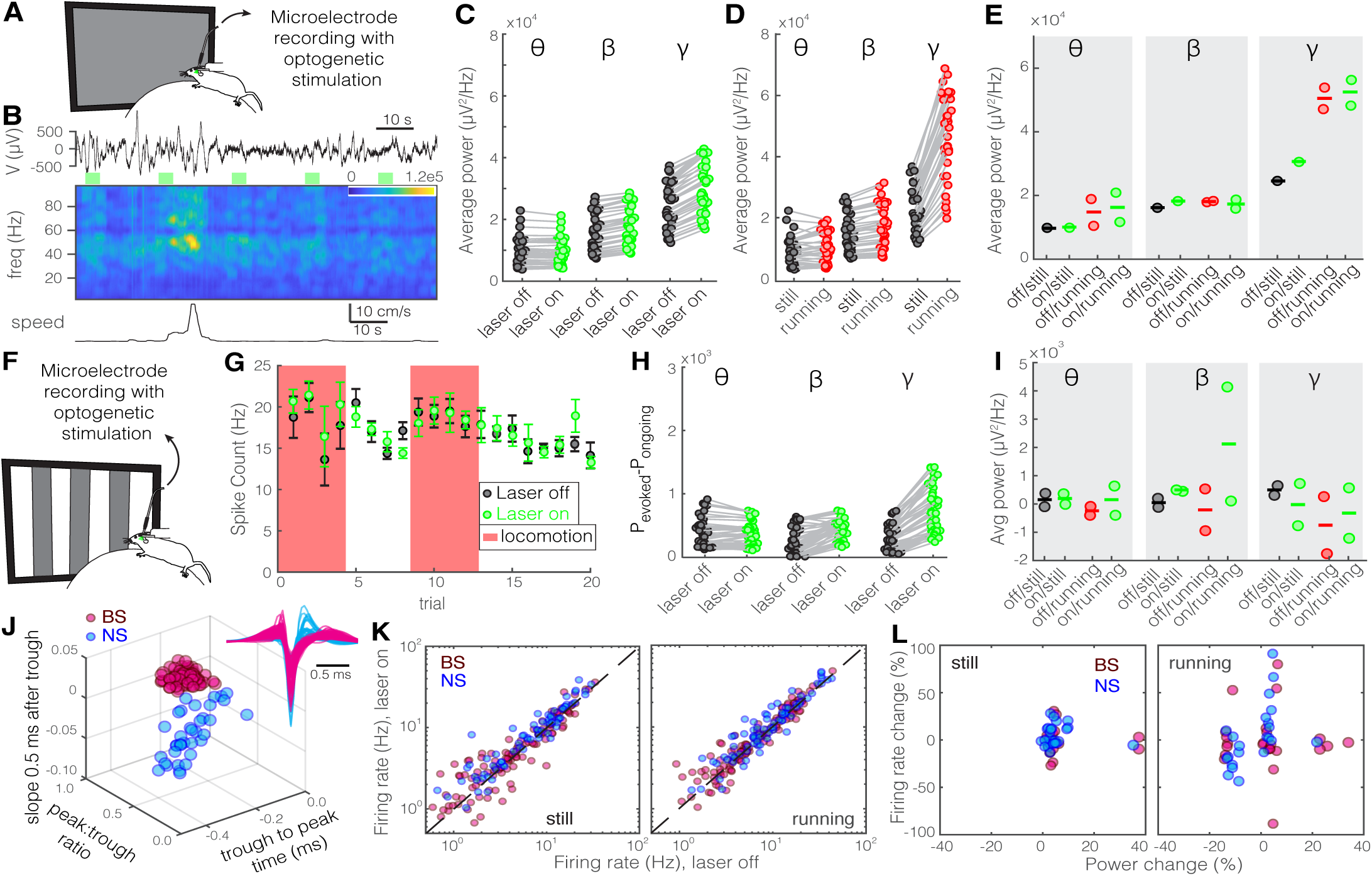
Optogenetic inhibition of PV nRT neurons does not significantly change gamma activity in V1 both with and without visual stimulation. Same format as Figure 2. Here are the statistics: **C**, theta: 9.8 vs. 9.8, p=5.1e-1, Wilcoxon-signed rank test; beta: 16.1 vs. 18.2, p=3.6e-7; gamma: 24.6 vs. 30.5, p=3.6e-7. **D**, theta: 9.8 vs. 10.4, p=0.07; beta: 16.1 vs. 17.9, p=4.0e-7; gamma: 24.6 vs. 47.1e3, p=3.6e-7. **E**, theta: 1.2e3 still & off vs. 1.2e3 still & on, p=1, 1.3e3 running & off vs. 2.0e3 running & on, p=0.34; beta: 1.6e3 still & off vs. 1.7e3 still & on, p=1, 1.7e3 running & off vs. 1.7e3 running & on, p=0.34; gamma: 2.5e3 still & off vs. 2.9e3 still & on, p=1, 5.3e3 running & off vs. 5.5e3 running & on, p=0.67. **H**, theta: 0.3e3 vs. 0.3e3, p=0.03; beta: 0.2e3 vs. 0.4e3, p=4.5e-9; gamma: 0.3e3 vs. 0.7e3, p=2.4e-9. K, BS & still: 6.7 (Hz) laser off vs. 7.0 laser on, p=1.8e-2; BS & running: 8.8 laser off vs. 9.4 laser on, p=1.0e-2; NS & still: 8.1 laser off vs. 9.5 laser on, p=1.0e-6; NS & running: 11.7 laser off vs. 13.1 laser on, p=1.9e-3.

**Figure 6–figure supplement 7.**
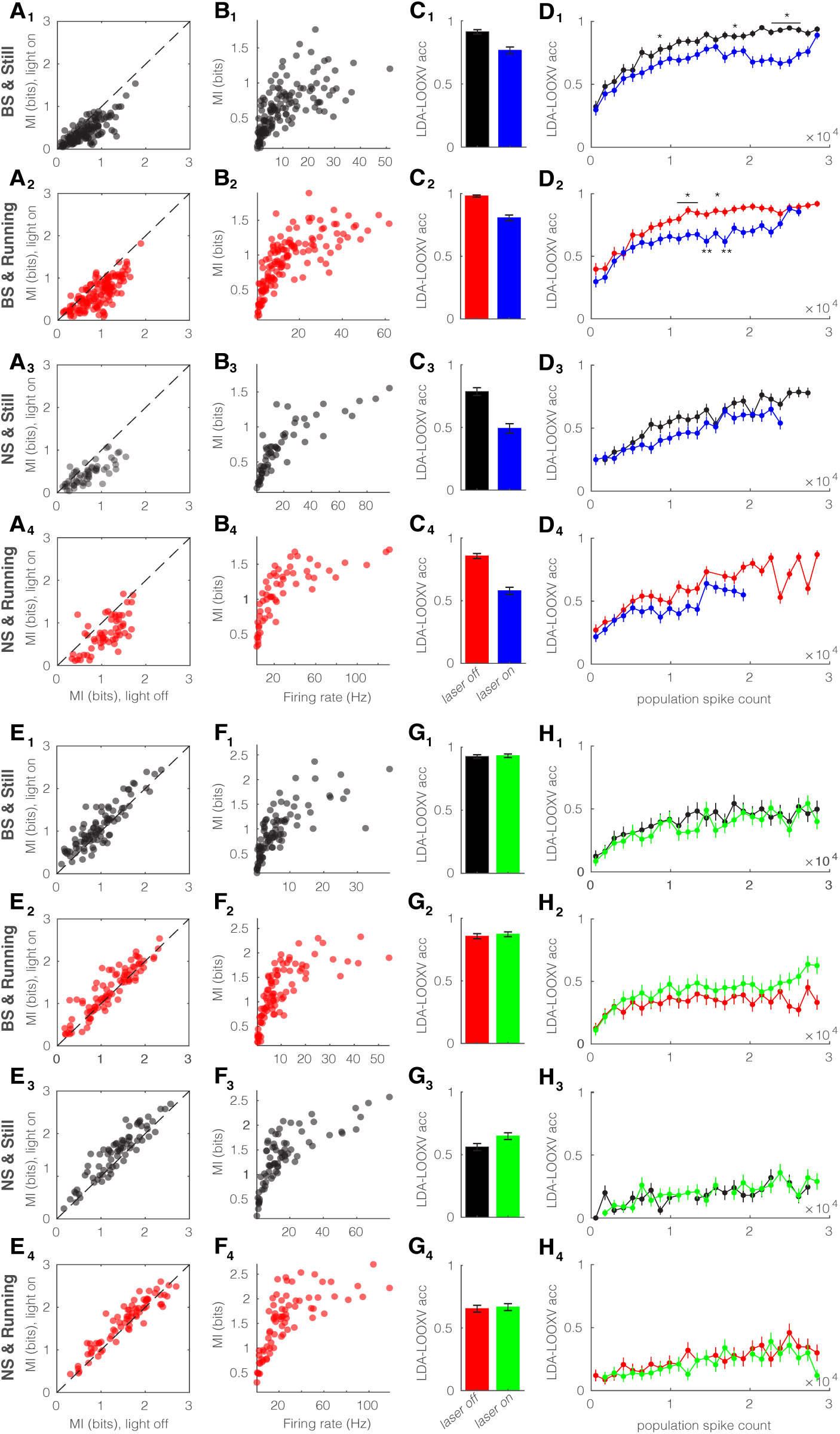
Optoge netic manipulation of SOM nRT neurons alters encoding accuracy in both BS and NS cells. **A**, Single-cell mutual information of BS neurons while stationary (A1) and running (A2), and NS neurons while stationary (A3) and running (A4) for optogenetic light-off versus light-on conditions (light-off to light-on; A1, 0.64 ± 0.03 to 0.42 ± 0.02, p=1e-8, n=141 cells, Wilcoxon signed-rank test; A2, 0.91 ± 0.03 to 0.64 ± 0.03, p=3e-9; A3, 0.69 ± 0.05 to 0.45 ± 0.03, p=3e-4, n=58; A4, 1.09 ± 0.05 to 0.76 ± 0.05, p=2e-5). Dashed line indicates unity. **B**, Single-cell mutual information against firing rate of BS neurons while stationary (B1) and running (B2), and NS neurons while stationary (B3) and running (B4) for light-off (rho, p, Spearman correlation; B1, 0.76, 7e-28; B2, 0.84, 4e-38; B3, 0.9, 0.0; B4, 0.86, 6e-18). **C**, Accuracy in LDA-LOOXV classification of visual stimulus movement orientation in BS neurons during stationary (C1) and running (C2), and NS neurons while stationary (C3) and running (C4) for light-off versus light-on (light-off to light-on; C1, 0.91 to 0.77, p=6e-6, Wilcoxon rank-sum test; C2, 0.98 to 0.81, p=2e-11; C3, 0.79 to 0.49, p=9e-9; C4, 0.86 to 0.58, p=1e-13). Error bars indicate boot-strapped estimates of SE. **D**, Classification accuracy for grating orientation as a function of population spike count for BS neurons during stationary (D1) and running (D2), and NS neurons during stationary (D3) and running (D4) for light-off versus light-on. Error bars indicate bootstrapped estimates of SE. Chance level would be at 0.16. *p<0.05, **p<0.01. **E**, Same as in A for optogenetic suppression of SOM cells in nRT (light-off to light-on; E1, 0.96 ± 0.05 to 1.10 ± 0.05, p=8e-7, n=90 cells; E2, 1.19 ± 0.06 to 1.31 ± 0.06, p=1e-4; E3, 1.35 ± 0.06 to 1.54 ± 0.07, p=9e-9, n=58; E4, 1.54 ± 0.07 to 1.65 ± 0.07, p=2e-5). **F**, Same as in B during optogenetic suppression (rho, p, Spearman correlation; F1, 0.84, 1e-24; F2, 0.84, 6e-25; F3, 0.81, 0.0; F4, 0.82, 2e-19). **G**, Same as in C during optogenetic suppression (light-off to light-on; G1, 0.93 to 0.93, p=0.75; G2, 0.86 to 0.87, p=0.53; G3, 0.56 to 0.65, p=0.03; G4, 0.65 to 0.67, p=0.71). **H**, Same as in D during optogenetic inhibition.

**Figure 6–figure supplement 8.**
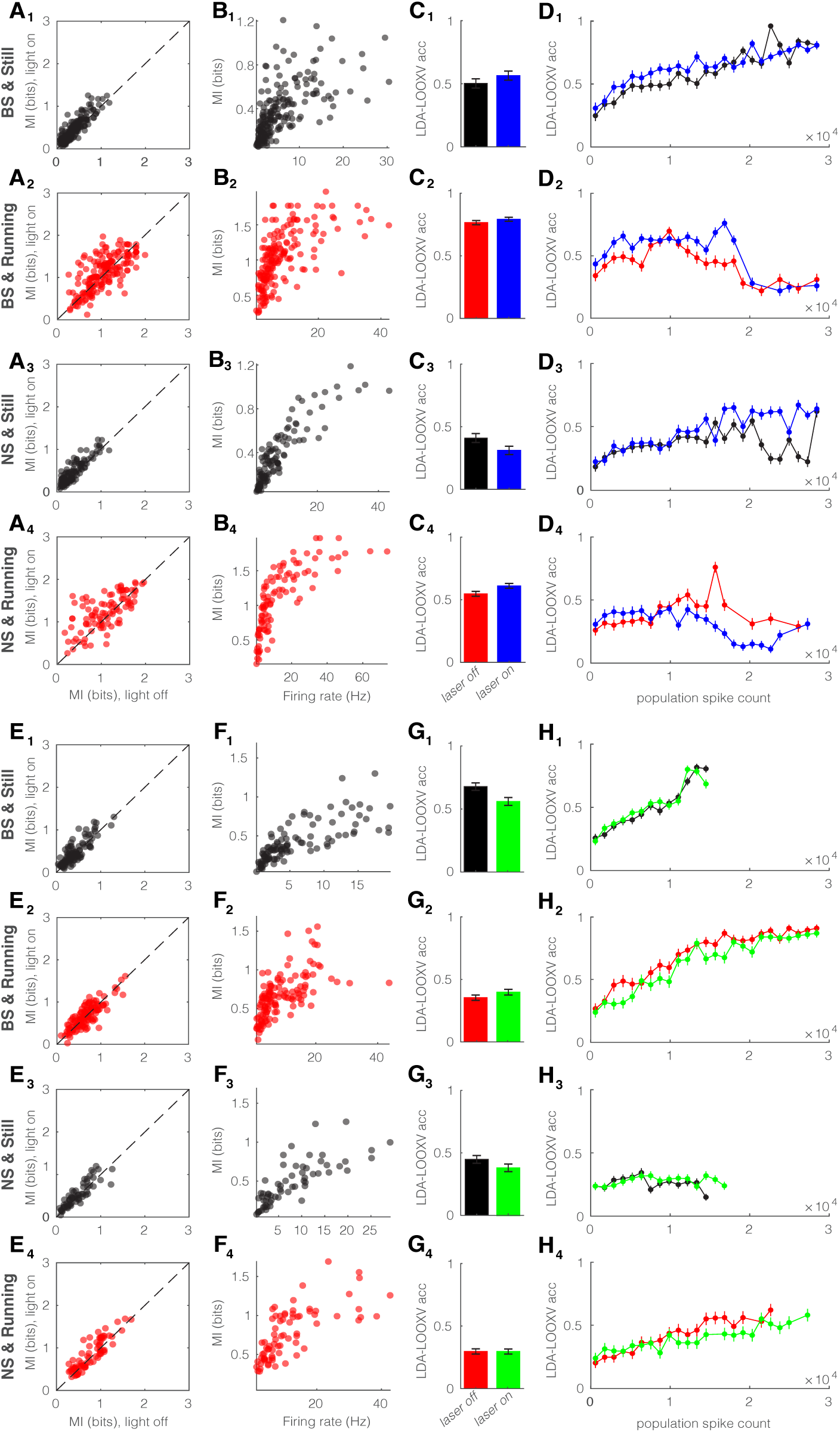
Optogenetic perturbation of PV nRT neurons does not alter encoding accuracy. Same format as Figure 6–figure supplement 6. Here are the statistics. **A**, Light-off to light-on; A1, 0.37 ± 0.02 to 0.44 ± 0.02, p=6e-12, n=191 cells, Wilcoxon signed-rank test; A2, 1.03 ± 0.03 to 1.08 ± 0.03, p=0.09; A3, 0.37 ± 0.02 to 0.45 ± 0.03, p=1.1e-8, n=113; A4, 1.09 ± 0.05 to 1.24 ± 0.04, p=2.5e-4. **B**, rho, p, Spearman correlation; B1, 0.74, 1.1e-32; B2, 0.72, 1.1e-29; B3, 0.89, 0.0; B4, 0.87, 1.1e-31. **C**, light-off to light-on; C1, 0.49 to 0.55, p=0.16, Wilcoxon rank-sum test; C2, 0.75 to 0.77, p=0.19; C3, 0.40 to 0.30, p=0.05; C4, 0.53 to 0.59, p=0.01. **E**, light-off to light-on; E1, 0.41 ± 0.02 to 0.46 ± 0.03, p=2.2e-3, n=127 cells; E2, 0.66 ± 0.02 to 0.67 ± 0.03, p=0.28; E3, 0.48 ± 0.04 to 0.52 ± 0.04, p=0.01, n=75; E4, 0.77 ± 0.04 to 0.84 ± 0.04, p=3.7e-4. **F**, rho, p, Spearman correlation; F1, 0.85, 2.3e-30; F2, 0.73, 4.5e-22; F3, 0.85, 4.0e-18; F4, 0.77, 0.0. **G**, Light-off to light-on; G1, 0.66 to 0.55, p=0.003; G2, 0.35 to 0.39, p=0.12; G3, 0.44 to 0.37, p=0.13; G4, 0.29 to 0.29, p=0.94.

**Figure 7–movie supplement 1. Optogenetic activation of SOM nRT neurons reduces the firing in dLGN neurons**.

**Figure 8–figure supplement 9.**
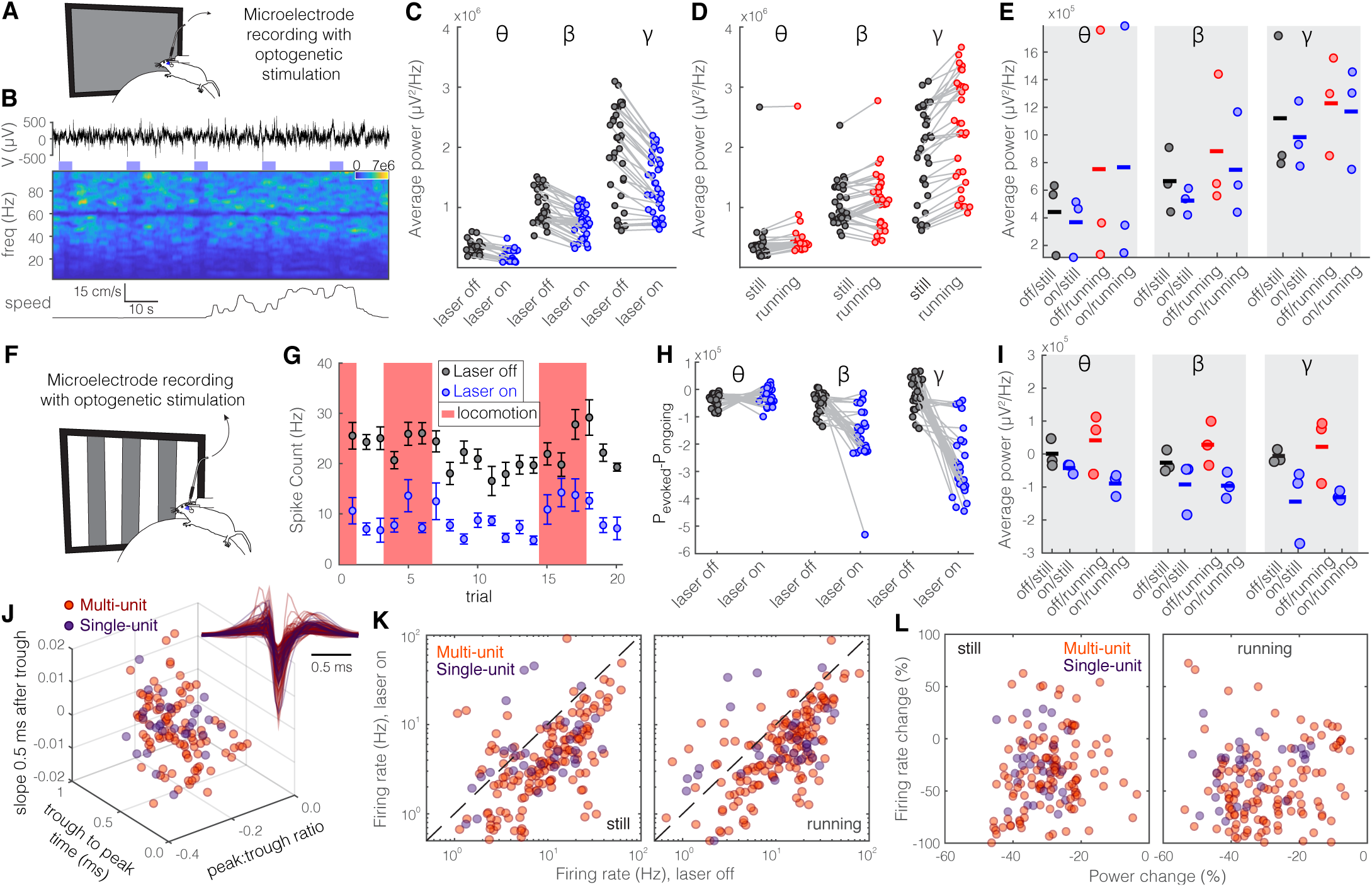
Optogenetic activation of SOM nRT neurons significantly reduces gamma activity in dLGN both with and without visual stimulation. Same format as Figure 2. Here are the statistics: **C**, theta: 146.0e3 vs. 133.4e3, p=1.1e-6, Wilcoxon-signed rank test; beta: 762.8e3 vs. 733.1e3, p=1.6e-5; gamma: 915.4e3 vs. 1098.3e3, p=5.4e-4. **E**, theta: 318.0e3 still & off vs. 371.5e3 still & on, p=0.70, 368.6e3 running & off vs. 442.6e3 running & on, p=0.40; beta: 1115.0e3 still & off vs. 1121.8e3 still & on, p=1, 884.3e3 running & off vs. 912.8e3 running & on, p=1; gamma: 2710.7e3 still & off vs. 2820.0e3 still & on, p=1, 2691.5e3 running & off vs. 2819.4e3 running & on, p=0.70. **J**, Height of the positive peak relative to the negative trough: MU −0.23 ± 0.02 vs. SU −0.25 ± 0.06 (p=0.98, Wilcoxon rank-sum test); the time from the negative trough to the peak: 0.59 ± 0.02 vs. 0.59 ± 0.04 ms (p=0.76); slope of the waveform 0.5 ms after the negative trough: 0.005 ± 0.001 vs. 0.002 ± 0.03 (p=0.69); (n=158 MU and n=35 SU cells). **K**, MU & still: 13.1 laser off vs. 6.2 laser on, p=4.4e-21; MU & running: 15.1 laser off vs. 8.0 laser on, p=1.6e-20; SU & still: 10.6 laser off vs. 7.5 laser on, p=0.008; SU & running: 14.4 laser off vs. 8.4 laser on, p=2.6e-3.

**Figure 8–figure supplement 10.**
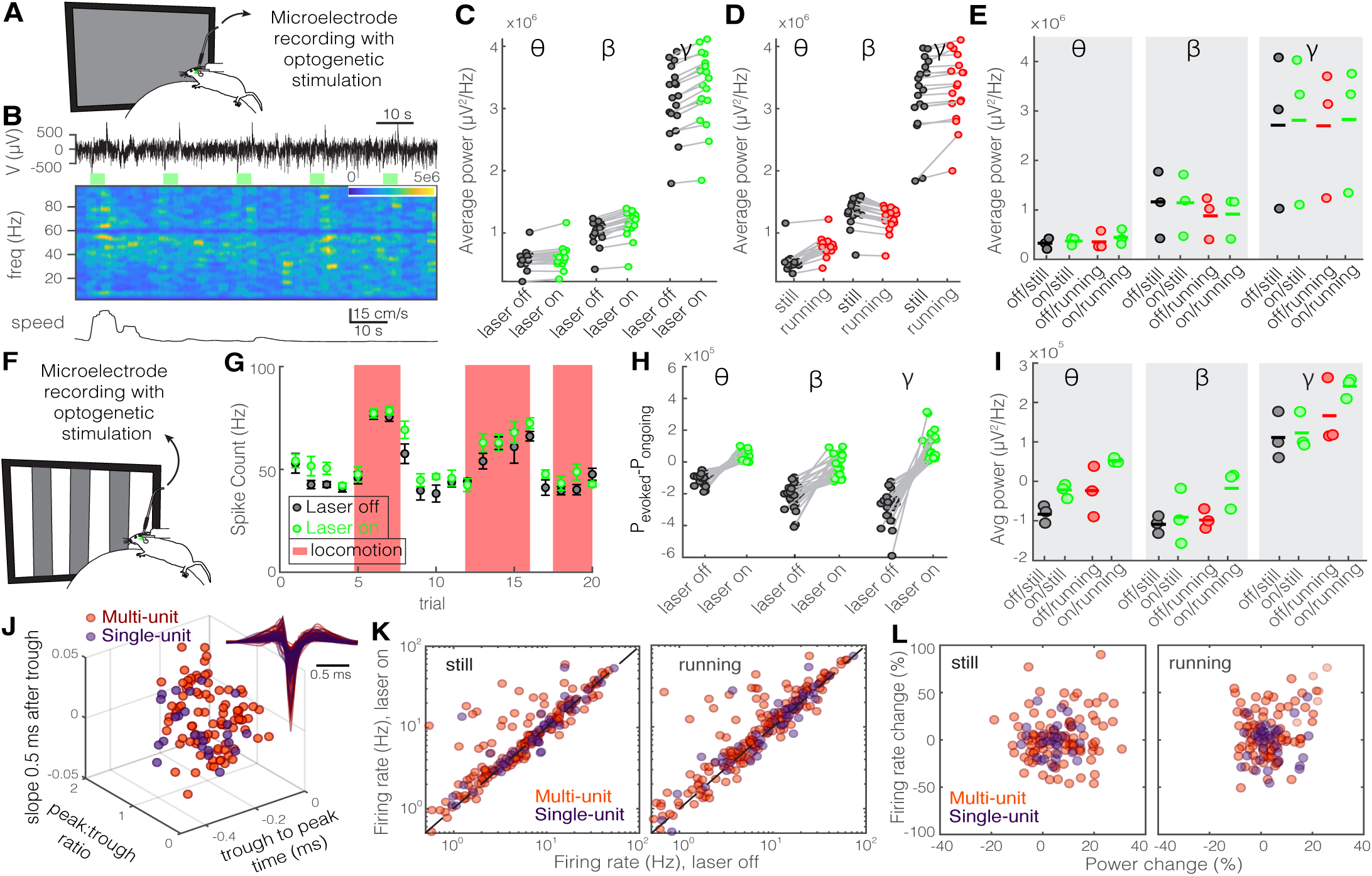
Optogenetic inhibition of SOM nRT neurons significantly enhances gamma ac tivity in dLGN both with and without visual stimulation. Same layout as Figure 2. Here are the statistics: **C**, theta: 575.2e3 vs. 609.5e3, p=3.2e-4, Wilcoxon-signed rank test; beta: 1021.0e3 vs. 1166.2e3, p=1.9e-4; gamma: 3137.6e3 vs. 3335.4e3, p=1.9e-4. **E**, theta: 442.1e3 still & off vs. 364.5e3 still & on, p=0.4, 752.4e3 running & off vs. 760.1e3 running & on, p=1; beta: 665.5e3 still & off vs. 134.8e3 still & on, p=0.4, 882.3e3 running & off vs. 748.3e3 running & on, p=0.7; gamma: 1120.1e3 still & off vs. 299.1e3 still & on, p=1, 1234.8e3 running & off vs. 1169.8e3 running & on, p=1. **J**, Height of the positive peak relative to the negative trough: MU −0.24 ± 0.01 vs. SU −0.22 ± 0.01 (p=0.24, Wilcoxon rank-sum test); slope of the waveform 0.5 ms after the negative trough: 0.57 ± 0.01 vs. 0.60 ± 0.03 ms (p=0.17); the time from the negative trough to the peak: 0.004 ± 0.001 vs. 0.003 ± 0.002 (p=0.64); (n=185 MU and n=49 SU cells). **K**, MU & still: 9.4 laser off vs. 11.3 laser on, p=1.1e-6; MU & running: 11.4 laser off vs. 14.2 laser on, p=6.3e-11; SU & still: 11.1 laser off vs. 12.3 laser on, p=0.14; SU & running: 13.8 laser off vs. 15.1 laser on, p=0.18.

## REFERENCES

Adaikkan C, Middleton SJ, Marco A, Pao P-C, Mathys H, Kim DN-W, Gao F, Young JZ, Suk H-J, Boyden ES, McHugh TJ, Tsai L-H (2019) Gamma Entrainment Binds Higher-Order Brain Regions and Offers Neuroprotection. Neuron 102:929–943.e8.

Ahlsén G, Lindström S, Lo F-S (1982) Functional distinction of perigeniculate and thalamic reticular neurons in the cat. Exp Brain Res 46:118–126.

Atencio CA, Schreiner CE (2008) Spectrotemporal Processing Differences between Auditory Cortical Fast-Spiking and Regular-Spiking Neurons. J Neurosci 28:3897–3910.

Barthó P, Hirase H, Monconduit L, Zugaro M, Harris KD, Buzsáki G (2004) Characterization of Neocortical Prin-cipal Cells and Interneurons by Network Interactions and Extracellular Features. J Neurophysiol 92:600–608.

Brainard DH (1997) The Psychophysics Toolbox. Spat Vis 10:433–436.

Buzsáki G, Wang X-J (2012) Mechanisms of Gamma Oscillations. Annu Rev Neurosci 35:203–225.

Campbell PW, Govindaiah G, Masterson SP, Bickford ME, Guido W (2020) Synaptic properties of the feedback connections from the thalamic reticular nucleus to the dorsal lateral geniculate nucleus. J Neurophysiol:jn.00757.2019.

Cardin J a, Carlén M, Meletis K, Knoblich U, Zhang F, Deisseroth K, Tsai L-H, Moore CI (2009) Driving fast-spiking cells induces gamma rhythm and controls sensory responses. Nature 459:663–667.

Chung JE, Magland JF, Barnett AH, Tolosa VM, Tooker AC, Lee KY, Shah KG, Felix SH, Frank LM, Greengard LF (2017) A Fully Automated Approach to Spike Sorting. Neuron 95:1381–1394.e6.

Clemente-Perez A, Makinson SR, Higashikubo B, Brovarney S, Cho FS, Urry A, Holden SS, Wimer M, Dávid C, Fenno LE, Acsády L, Deisseroth K, Paz JT (2017) Distinct Thalamic Reticular Cell Types Differentially Modulate Normal and Pathological Cortical Rhythms. Cell Rep 19:2130–2142.

Crabtree JW (2018) Functional Diversity of Thalamic Reticular Subnetworks. Front Syst Neurosci 12.

Crick F (1984) Function of the thalamic reticular complex: The searchlight hypothesis. Proc Natl Acad Sci U S A 81:4586–4590.

Dadarlat MC, Stryker MP (2017) Locomotion Enhances Neural Encoding of Visual Stimuli in Mouse V1. J Neurosci 37:3764–3775.

Doesburg SM, Roggeveen AB, Kitajo K, Ward LM (2008) Large-Scale Gamma-Band Phase Synchronization and Selective Attention. Cereb Cortex 18:386–396.

Du J, Blanche TJ, Harrison RR, Lester HA, Masmanidis SC (2011 Multiplexed, high density electrophysiology with nanofabricated neural probes. PLoS One 6.

Engel AK, Fries P, Singer W (2001) Dynamic predictions: Oscillations and synchrony in top–down processing. Nat Rev Neurosci 2:704–716.

Fu Y, Kaneko M, Tang Y, Alvarez-Buylla A, Stryker MP (2015) A cortical disinhibitory circuit for enhancing adult plasticity. Elife 4:1–12.

Fu Y, Tucciarone JM, Espinosa S, Sheng N, Darcy DP, Nicoll RA, Huang ZJ, Stryker MP (2014) A Cortical Circuit for Gain Control by Behavioral State. Cell 156:1139–1152.

Gentet LJ, Ulrich D (2003) Strong, reliable and precise synaptic connections between thalamic relay cells and neurones of the nucleus reticularis in juvenile rats. J Physiol 546:801–811.

Gray CM, Engel AK, König P, Singer W (1992) Synchronization of oscillatory neuronal responses in cat striate cortex: Temporal properties. Vis Neurosci 8:337–347.

Halassa MM, Acsády L (2016) Thalamic Inhibition: Diverse Sources, Diverse Scales. Trends Neurosci 39:680–693.

Hoseini MS, Rakela B, Flores-Ramirez Q, Hasenstaub AR, Alvarez-Buylla A, Stryker MP (2019) Transplanted Cells Are Essential for the Induction But Not the Expression of Cortical Plasticity. J Neurosci 39:7529–7538.

Houser CR, Vaughn JE, Barber RP, Roberts E (1980) GABA neurons are the major cell type of the nucleus reticularis thalami. Brain Res 200:341–354.

Kaneko M, Fu Y, Stryker MP (2017) Locomotion Induces Stimulus-Specific Response Enhancement in Adult Visual Cortex. J Neurosci 37:3532–3543.

Kaneko M, Stryker MP (2014) Sensory experience during locomotion promotes recovery of function in adult visual cortex. Elife 2014:1–16.

Kleiner M, Brainard DH, Pelli DG (2007) What’s new in Psychtoobox-3? Perception 36:14.

Kreiter A, Singer W (1996) Stimulus-dependent synchronization of neuronal responses in the visual cortex of the awake macaque monkey. J Neurosci 16:2381–2396.

Lam Y-W, Sherman SM (2011) Functional Organization of the Thalamic Input to the Thalamic Reticular Nucleus. J Neurosci 31:6791–6799.

Lee AM, Hoy JL, Bonci A, Wilbrecht L, Stryker MP, Niell CM (2014) Identification of a Brainstem Circuit Regulating Visual Cortical State in Parallel with Locomotion. Neuron 83:455–466.

Lein ES et al. (2007) Genome-wide atlas of gene expression in the adult mouse brain. Nature 445:168–176.

McAlonan K, Cavanaugh J, Wurtz RH (2008) Guarding the gateway to cortex with attention in visual thalamus. Nature 456:391–394.

McCormick DA, Bal T (1997) SLEEP AND AROUSAL: Thalamocortical Mechanisms. Annu Rev Neurosci 20:185–215.

Moore AK, Weible AP, Balmer TS, Trussell LO, Wehr M (2018) Rapid Rebalancing of Excitation and Inhibition by Cortical Circuitry. Neuron 97:1341–1355.e6.

Niell CM, Stryker MP (2008) Highly Selective Receptive Fields in Mouse Visual Cortex. J Neurosci 28:7520–7536.

Niell CM, Stryker MP (2010) Modulation of Visual Responses by Behavioral State in Mouse Visual Cortex. Neuron 65:472–479.

Pavlova M, Birbaumer N, Sokolov A (2006) Attentional Modulation of Cortical Neuromagnetic Gamma Response to Biological Movement. Cereb Cortex 16:321–327.

Paxinos G, Franklin KBJ (2001) Paxinos and Franklin’s the Mouse Brain in Stereotaxic Coordinates.

Paz JT, Bryant AS, Pend K, Fenno L, Yizhar O, Frankel WN, Deisseroth K, Huguenard JR (2011) A new mode of corticothalamic transmission revealed in the Gria4 model of absence epilepsy. Nat Neurosci 14: 1167:1173

Paz JT, Davidson TJ, Frechette ES, Delord B, Parada I, Peng K, Deisseroth K, Huguenard JR (2013) Closed-loop optogenetic control of thalamus as a tool for interrupting seizures after cortical injury. Nat Neurosci 16:64–70.

Phillips EA, Hasenstaub AR (2016) Asymmetric effects of activating and inactivating cortical interneurons. Elife 5:e18383.

Piscopo DM, El-Danaf RN, Huberman AD, Niell CM (2013) Diverse Visual Features Encoded in Mouse Lateral Geniculate Nucleus. J Neurosci 33:4642–4656.

Quiroga RQ, Reddy L, Kreiman G, Koch C, Fried I (2005) Invariant visual representation by single neurons in the human brain. Nature 435:1102–1107.

Ray S, Crone NE, Niebur E, Franaszczuk PJ, Hsiao SS (2008a) Neural Correlates of High-Gamma Oscillations (60-200 Hz) in Macaque Local Field Potentials and Their Potential Implications in Electrocorticography. J Neurosci 28:11526–11536.

Ray S, Niebur E, Hsiao SS, Sinai A, Crone NE (2008b) High-frequency gamma activity (80–150Hz) is increased in human cortex during selective attention. Clin Neurophysiol 119:116–133.

Reinhold K, Lien AD, Scanziani M (2015) Distinct recurrent versus afferent dynamics in cortical visual processing. Nat Neurosci 18:1789–1797.

Ritter-Makinson S, Clemente-Perez A, Higashikubo B, Cho FS, Holden SS, Bennett E, Chkhaidze A, Eelkman Rooda OHJ, Cornet M-C, Hoebeek FE, Yamakawa K, Cilio MR, Delord B, Paz JT (2019) Augmented Reticular Thalamic Bursting and Seizures in Scn1a-Dravet Syndrome. Cell Rep 26:54–64.e6.

Saleem AB, Lien AD, Krumin M, Haider B, Rosón MR, Ayaz A, Reinhold K, Busse L, Carandini M, Harris KD, Carandini M (2017) Subcortical Source and Modulation of the Narrowband Gamma Oscillation in Mouse Visual Cortex. Neuron 93:315–322.

Sherman SM, Koch C (1986) The control of retinogeniculate transmission in the mammalian lateral geniculate nucleus. Exp Brain Res 63:1–20.

Siegel M, Donner TH, Oostenveld R, Fries P, Engel AK (2008) Neuronal Synchronization along the Dorsal Visual Pathway Reflects the Focus of Spatial Attention. Neuron 60:709–719.

Singer W, Gray CM (1995) Visual feature integration and the temporal correlation hypothesis. Annu Rev Neurosci 18:555–586.

Sohal VS, Zhang F, Yizhar O, Deisseroth K (2009) Parvalbumin neurons and gamma rhythms enhance cortical circuit performance. Nature 459:698–702.

Stryker MP (1989) Is grandmother an oscillation? Nature 338:297–298.

Taylor K, Mandon S, Freiwald WA, Kreiter AK (2005) Coherent Oscillatory Activity in Monkey Area V4 Predicts Success ful Allocation of Attention. Cereb Cortex 15:1424–1437.

Wang X (2010) Neurophysiological and computational principles of cortical rhythms in cognition. Physiol Rev 90:1195–1268

Wells MF, Wimmer RD, Schmitt LI, Feng G, Halassa MM (2016) Thalamic reticular impairment underlies attention deficit in Ptchd1Y mice. Mature 532:58–63.

Yazdan-Shahmorad A, Kipke DR, Lehmkuhle MJ (2013) High gamma power in ECoG reflects cortical electrical stimulation effects on unit activity in layers V/VI. J Neural Eng 10:066002.

Zikopoulos B, Barbas H (2012) Pathways for Emotions and Attention Converge on the Thalamic Reticular Nucleus in Primates. J Neurosci 32:5338–5350.

